# Decoding the transcriptome of denervated muscle at single-nucleus resolution

**DOI:** 10.1101/2021.10.25.463678

**Authors:** Hongchun Lin, Xinxin Ma, Yuxiang Sun, Hui Peng, Yanlin Wang, Sandhya Sara Thomas, Zhaoyong Hu

**Author notes:** These authors contributed equally to this work (Hongchun Lin, Xinxin Ma). Corresponding author: Zhaoyong Hu, M.D., Nephrology Division, Department of Medicine, Baylor College of Medicine One Baylor Plaza, ABBR R702, Houston, TX 77030, USA.

## Abstract

**Background:** Skeletal muscle exhibits remarkable plasticity under both physiological and pathological conditions. One major manifestation of this plasticity is muscle atrophy that is an adaptive response to catabolic stimuli. Since the heterogeneous transcriptome responses to catabolism in different types of muscle cells are not fully characterized, we applied single-nucleus RNA sequencing (snRNA-seq) to unveil muscle atrophy related transcriptional changes at single nucleus resolution.

**Methods:** Using a sciatic denervation mouse model of muscle atrophy, snRNA-seq was performed to generate single-nucleus transcriptional profiles of the gastrocnemius muscle from normal and denervated mice. Various bioinformatics analyses, including unsupervised clustering, functional enrichment analysis, trajectory analysis, regulon inference, metabolic signature characterization and cell-cell communication prediction, were applied to illustrate the transcriptome changes of the individual cell types.

**Results:** A total of 29,539 muscle nuclei (normal *vs* denervation: 15,739 *vs* 13, 800) were classified into 13 nuclear types according to the known cell markers. Among these, the type IIb myonuclei were further divided into two subgroups, which we designated as type IIb1 and type IIb2 myonuclei. In response to denervation, the proportion of type IIb2 myonuclei increased sharply (78.12% *vs* 38.45%, *p* <0.05). Concomitantly, trajectory analysis revealed that denervated type IIb2 myonuclei clearly deviated away from the normal type IIb2 myonuclei, indicating that this subgroup underwent robust transcriptional reprogramming upon denervation. Signature genes in denervated type IIb2 myonuclei included *Runx1*, *Gadd45a*, *Igfn1*, *Robo2*, *Dlg2*, and *Sh3d19* (*p* <0.001). The gene regulatory network analysis captured a group of atrophy-related regulons (Foxo3, Runx1, Elk4, Bhlhe40) whose activities were enhanced (*p* <0.01), especially in the type IIb2 myonuclei. The metabolic landscape in the myonuclei showed that most of the metabolic pathways were downregulated by denervation (*p* <0.001), while some of the metabolic signaling, such as glutathione metabolism, was specifically activated in the denervated type IIb2 myonulei. We also investigated the transcriptomic alterations in the type I myofibers, muscle stem cells, fibro-adipogenic progenitors, macrophages, endothelial cells and pericytes and characterized their signature responses to denervation. By predicting the cell-cell interactions, we observed that the communications between myofibers and muscle resident cells were diminished by denervation.

**Conclusion:** Our results define the myonuclear transition, metabolic remodeling and gene regulation networks reprogramming associated with denervation-induced muscle atrophy and illustrate the molecular basis of the heterogeneity and plasticity of muscle cells in response to catabolism. These results provide a useful resource for exploring the molecular mechanism of muscle atrophy.

## Introduction

Skeletal muscle atrophy occurs in response to a variety of clinical conditions, including cachexia, chronic kidney disease, and denervation. It is characterized by a loss of muscle mass and contractile function, which contributes to reduced quality of life of patients [1], and in some circumstances increased mortality. Therefore, it is paramount to understand the transcriptional landscape of atrophic muscle and explore the molecular mechanisms behind muscle atrophy, in order to develop corresponding treatment strategies.

Myofibers are giant muscle cells with multiple myonuclei and are typically classified as type I and type II fibers. In mice, the type II myofiber can be further divided into type IIa, IIx, and type IIb [2]. Meanwhile, the interstitial spaces between such fibers are populated with muscle-resident cells, including fibro-adipogenic progenitors (FAPs), vascular cells, nerve cells, and immune cells such as macrophages and lymphocytes. The exquisite orchestration of these heterogeneous cell populations is required for the maintenance of muscle function and homeostasis. To this end, distinct gene regulatory networks initiate diverse gene expression patterns in each type of muscle cell, leading the tissue to feature complex transcriptional heterogeneity. In this context, examining bulk gene expression may not allow the delineation of transcriptional profiles in a cell-specific manner.

Single-cell RNA sequencing can simultaneously interrogate gene expression and signaling pathways in multiple cell types within a tissue. It has been used to characterize the gene expression profiles of normal or unperturbed muscle tissues in both mouse and human[3, 4]; in particular, Puri and colleagues precisely described the response of muscle-resident cells to denervation at single-cell resolution[5]. With advances in library preparation and isolation techniques, single-nucleus RNA sequencing (snRNA-seq) has enabled the detection of transcriptional states in multinucleated cells such as myofibers; for example, Dos Santos et al. ingeniously defined the coordinated regulation of multiple-nuclear transcription in myofibers using snRNA-seq [6]. In another elegant study, Petrany et al. depicted the transcriptional heterogeneity of normal myofibers and identified new myonuclear types [7]. However, while these studies provide integral parts that have allowed us to build complete gene expression landscapes for normal muscles or perturbed muscle-resident cells, the transcriptional heterogeneity of myofibers in response to catabolic stimuli remains to be established.

In this study, we employed snRNA-seq to elucidate transcriptional heterogeneity in denervated myofibers. We chose the denervation context because it induces pronounced muscle atrophy and is relevant to clinical conditions including neurodegenerative diseases and age-related muscle atrophy and weakness [8]. Importantly, the model is standardized and reproducible, which is less likely to be affected by variations in disease progression and other complications such as anorexia or systemic inflammation. In addition, denervation-induced muscle atrophy is initiated by the loss of neural signals at myofibers. As such, studying this phenotype specifically facilitates the prediction of communications between myofibers and other non-parenchymal cells during muscle atrophy. Using this model, we identified unique atrophic-responsive gene signatures of all muscle tissue populations and revealed uncharacterized diversity within denervated myofibers and other resident cells. Further, we demonstrated the specific gene regulation networks (regulons) that govern the expression of those gene signatures, thus, providing a comprehensive single-nucleus/cell atlas of atrophic muscle.

## Methods

### Mouse model and sample collection

12-week-old, male C57BL/6J mice (The Jackson Laboratory, Bar Harbor, ME) (n=3) were used as sciatic nerve transection model. Procedures of denervation surgery were carried out as previously described [9]. Briefly, mice were anesthetized (5% isoflurane) and the left sciatic nerve was removed for a 2-mm piece. Gastrocnemius (Gas) muscles were harvested 2 weeks after denervation. Gas muscles from left hind limb of normal mice served as controls (n=3). All mice were housed under specific pathogen free condition, with a 12/12 h light/dark cycle. All procedures were approved by the Baylor College of Medicine Institutional Animal Care and Use Committee and carried out in accordance with the recommendations of Institutional and NIH guidelines.

### Nuclei isolation from skeletal muscles

The muscle samples were processed according to a published nuclear isolation protocol [10] with modifications. We first prepared nuclei isolation media1 (NIM1: 320 mM sucrose, 25mM KCl, 5 mM MgCl2, and 10 mM Tris buffer, pH=8.0) and homogenization buffer [NIM1 buffer containing 1 µM DTT, 0.4 U/μL RNase Inhibitor (NEB Inc, Ipswich, MA), 0.20U/μL SUPERase-In RNase Inhibitor (Thermo Fisher scientific Inc, Waltham, MA), and 0.1% Triton X-100]. Then, Gas muscles (100∼150 mg) were minced on ice with a razor blade. Muscles were resuspended in 5 mL ice-cold homogenization buffer and homogenized with a dounce grinder for 12 strokes gently on ice. The homogenate was sequentially filtered through 70 mm and 40 mm cell strainers (pluriSelect, El Cajon, CA) and centrifuged (x1000g) for 5 minutes at 4°C to pellet the nuclei. The pellet was resuspended with 1 mL of ice-cold wash buffer (PBS containing 2% bovine serum albumin and 0.2U/μL ribonuclease inhibitor) and then filtered through the 20μm cell strainer before centrifugation (x500g) for 5 minutes at 4°C. The collected nuclei were resuspended in 100 μL of cold wash buffer and the purity of nuclei suspensions was confirmed by microscopy with DAPI staining. All chemicals were purchased from Sigma-Aldrich Chemicals (St. Louis, MO) unless otherwise noted.

### Library preparation for single-nucleus RNA-sequencing (snRNA-seq)

Nuclei suspensions were adjusted to a concentration of 1200 nuclei/μL using a hemocytometer and were loaded into the 10x Chromium Chip (∼ 10,000 nuclei per sample). snRNA-seq libraries were prepared using Chromium Single Cell 3’ Reagent Kits v3 and sequenced on the Illumina HiSeq4000 platform using a custom paired-end sequencing mode as previously described [11].

### snRNA-seq data processing and analysis

Raw sequencing data were processed using Cell Ranger Single Cell Software Suite (v3.0.2) (10x Genomics). A custom reference (mm10 pre-mRNA) was built for reads alignment in accordance with 10x Genomics recommendations and all reads were mapped to the reference genome using STAR (Spliced Transcripts Alignment to a Reference) with default setting. The mapped reads with valid barcodes and unique molecular identifiers (UMIs) were used to generate the gene-barcode matrix. Further analysis was performed using Seurat package (v3.1.0) in R (v3.6.1) [12]. A quality control step was performed by removing doublets (DoubletFinder v2.0) [13] and low-quality nuclei (less than 200 genes or more than 0.8% mitochondria genes) (Seurat v3.1.0) to ensure only transcripts originating from nuclei were retained. Genes detected in less than 3 nuclei were also removed. The resulting expression matrix of each sample was normalized using the NormalizeData function. Then, the IntegrateData function was applied to integrate datasets and ScaleData function was used to scale the data.

1. ***Dimension reduction and clustering*:** The top 2000 variable genes were used for downstream principal component analysis (PCA) prior to dimensionality reduction. Top 10 principal components were then input for Uniform Manifold Approximation and Projection (UMAP) dimensionality reduction. Clusters were identified using FindCluster function (resolution = 0.5). The cluster annotation was based on the expression of established cell type identities and refined using the signature genes identified by FindAllMarkers function (Wilcoxon Rank Sum test, min.pct = 0.25, logfc.threshold = 0.25).
2. ***Differential expressed genes (DEGs) analysis:*** Differential expression analysis was performed using Seurat’s FindMarkers function (Wilcoxon Rank Sum test, min.pct = 0.25, logfc.threshold = 0.25).
3. ***Kyoto Encyclopedia of Genes and Genomes (KEGG) pathway enrichment analysis:*** Pathways analysis on the differential expressed genes was performed by using the enrichKEGG function in the ClusterProfile package (version 3.12.0) [14]. A *p* value <0.05 and an adjusted p value < 0.05 was considered statistically significant.
4. ***Gene set enrichment analysis (GSEA)*** GSEA analysis was conducted using the javaGSEA software (v4.1.0) (http://software.broadinstitute.org/gsea/downloads.jsp.). Normalized read counts for genes from normal and denervation samples were input for analysis. Gene sets with enrichment scores of *p* value <0.05 and false discovery rate (FDR) < 0.25 were considered significantly enriched.

### Trajectory analysis

Trajectory analysis was performed using the Monocle3 (version 0.2.3) algorithm [15] with default parameters. The normalized counts matrix for the whole dataset or each cell/nucleus type were input to create CDS objects, followed by data normalization and PCA analysis, dimension reduction and cell clustering (cluster_cells, resolution=0.0001). After principal graph was plotted, the nulcei were ordered in pseudotime. All the trajectory graphs were visualized.

### Prediction of gene regulatory networks (GRNs) using SCENIC

The SCENIC (version 1.1.2) [16] was applied to predict the potential GRNs regulating normal and denervated muscles. We first inferred the regulons that consist of upstream transcription factor (TF) and its candidate downstream target genes using GENIE3. We analyzed these regulons with RcisTarget. Target genes exhibiting significant motif enrichment of corresponding TF were retained, while indirect target genes without motif enrichment were removed. The genomic regions for TF-motif search were limited to 10kb around the transcription start site (genome reference mm10). Next, we scored the activity of each regulon in each nucleus with AUCell to identify the active regulons in different cell/nucleus types (such as myonuclei, MuSCs, FAPs and macrophages), which were further visualized using pheatmap R package. We also performed a clustering on the resulting binary regulon activity matrix to assess the different of cellular states between normal and denervation, and visualized the active regulons driving the relevant states in t-SNE plot.

### Evaluation of metabolic pathway activity

The pipeline Single-Cell-Metabolic-Landscape [17] was used to characterize the metabolic features of different myonuclei subtypes and the metabolic changes in response to denervation. This approach evaluates the activity of metabolic pathway based on an activity score defined as relative gene expression value averaged over all genes in one metabolic pathway. Metabolic genes (1566) and pathways (85) were obtained from the KEGG database. The metabolic genes identified in myonuclei of normal and denervated muscles were input for UMAP (log-transformed gene counts) to visualize the effect of metabolic genes on clustering and cell states. The scores of pathways activity in myonuclei subtypes were calculated and visualized. Score of pathway activity with *p* value < 0.01 (random permutation test) was considered statistically significant. Outlier genes with relative expression levels greater than 3 times 75th percentile or less than 1/3 times 25th percentile in each pathway were excluded.

### Prediction of ligand-receptor (L-R) interaction

We applied a ligand-receptor interaction model [3] to predict the intercellular communication mediating by ligand-receptor interactions between different muscle cells. An integrated database from NicheNet package [18] was used for L-R pair references. We prioritized ligands that are differentially expressed (logfc>0.25, min.pct=0.25) in type I/II myofibers compared with other cell types, and their differentially expressed receptors. We calculated the score for a potential L-R pair by multiplying the average ligand expression in myofibers and the average receptor expression in other cell type.

### Statistical analysis

All statistical computing were performed using the R software. Unless otherwise specified, comparisons between two groups were performed using Student’s t-test or Wilcoxon rank-sum test. The level of significance was set at \**p* < 0.05, \*\**p* < 0.01, or \*\*\**p* < 0.001.

### Data Availability

Raw and processed sequencing data have been deposited in GEO (GSE183802) (https://www.ncbi.nlm.nih.gov/geo/query/acc.cgi?acc=GSE183802).

## Results

### Muscle heterogeneity and cell type composition in normal and denervated muscles

To understand how muscle cell heterogeneity responds to denervation, we created a sciatic denervation model using 12-week-old male C57BL/6J mice and harvested gastrocnemius (Gas) muscle at two weeks post-denervation. Denervation caused severe atrophy in both type I and type II myofibers (Figure 1A and B, Figure S1A) and led to the fiber size distribution shifting obviously left-toward (Figure 1C). Purified nuclei from Gas muscle cells were used to construct single-nucleus libraries with the 10x Genomics Chromium system, which were sequenced as previously described [11]. After quality control, a total of 29,539 nuclei were retained, comprising 15,739 nuclei from three normal mice and 13,800 nuclei from three denervated mice. Using the Seurat package (3.1.0), we identified 13 nuclear types representing nine major cell types in muscles: type I myofibers, type II myofibers, muscle stem cells (MuSCs, satellite cells), FAPs, macrophages, endothelial cells, pericytes, and other cell types (Figure 1D and Figure S1B). Cell identities and their top five signature genes are provided in Figure 1E and Figure S1C.

**Figure 1.**
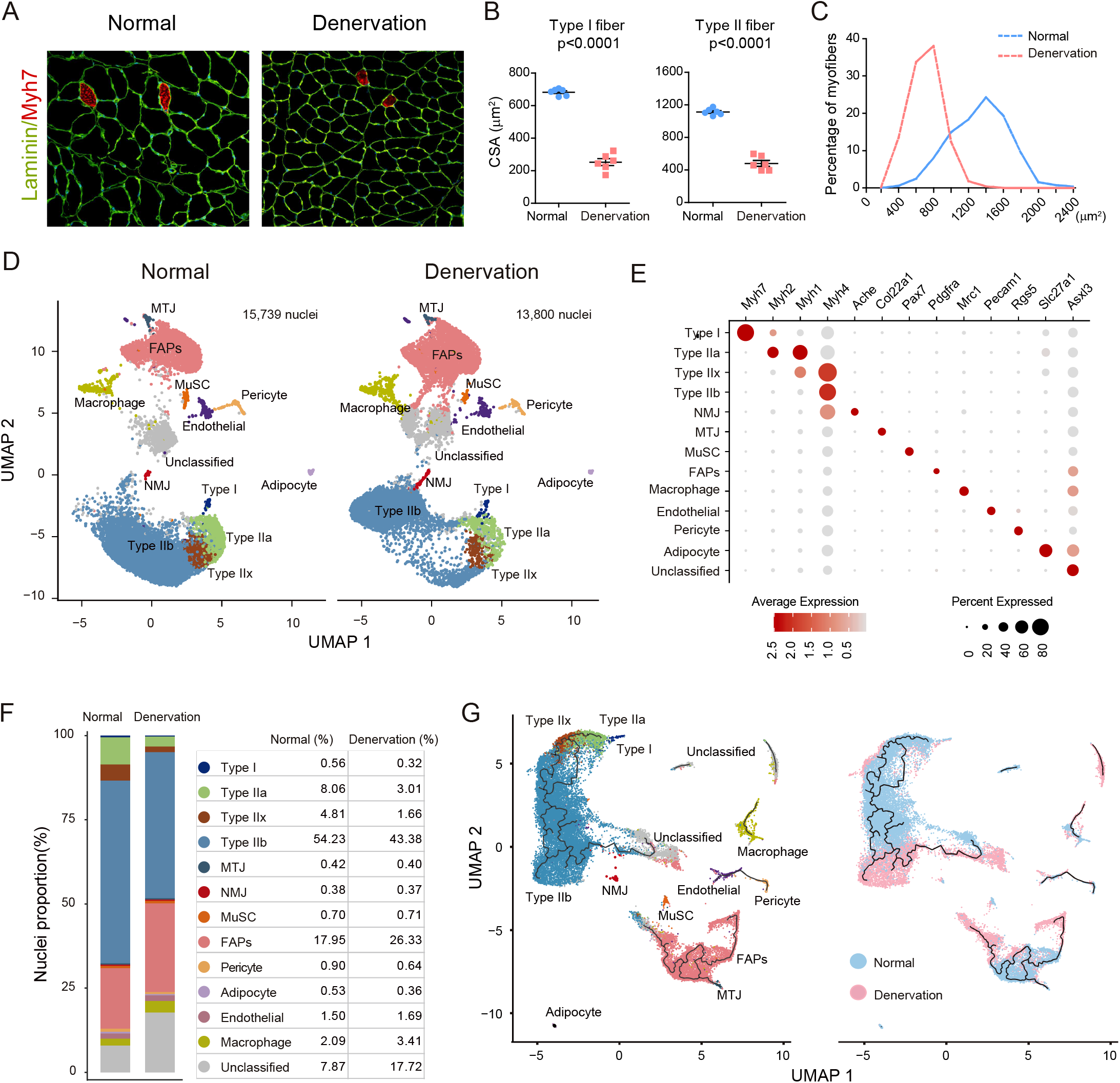
Classification of nucleus/cell types in normal and denervated gastrocnemius (Gas) muscles. (A) Immunofluorescent staining: Type I fibers are labeled with anti-Myh7 antibody (red) and co-stained with laminin (green). Images were taken at ×20 magnification. (B) Comparison of cross-sectional area (CSA) between normal and denervated conditions. The Data are presented as mean ± SEM (n=6). (C) Fiber size distribution of normal and denervated Gas muscles. The size of 300-400 fibers in muscles of each animal was assessed (n=6). (D)Uniform Manifold Approximation and Projection (UMAP) visualized nuclear clusters, which are colored and labeled according to cell identities, in normal (left panel) and denervated (right panel) Gas muscles. Type I: type I myonuclei; Type IIa: type IIa myonuclei; Type IIx: type IIx myonuclei; Type IIb: type IIb myonuclei; MTJ: myotendinous junction nuclei; NMJ: neuromuscular junction nuclei; MuSCs: muscle satellite cells nuclei; FAPs: fibro-adipogenic progenitors’ nuclei; macrophage: macrophages nuclei; ECs: endothelial cells nuclei; Pericytes: pericytes nuclei; Adipocyte: adipocyte nuclei. (E) Dot plot displaying the cell identity of each nuclear cluster. (F) Proportion of nuclear types in normal and denervated muscles. Each nucleus/cell type is color-coded. (G) UMAP plots with pseudotime trajectories of all nuclei obtained from normal and denervated muscles. The black lines on the UMAP plots represent branched trajectories. Each point denotes a single nucleus. Left panel: nuclei are color-coded according to their cluster assignments in (D). Right panel: nuclei are colored by conditions (blue: normal; pink: denervation).

We first analyzed the changes of cell populations in muscles following denervation. In normal gastrocnemius muscle, type II myonuclei was the predominant cell type (67% of total examined nuclei); type I myonuclei and satellite cells only accounted for 0.56% and 0.7% of cells, respectively, while non-myofiber nuclei such as FAPs (17.95%), macrophages (2.09%), and endothelial cells (1.5%) were frequently detected. These results are comparable with previous reports [6, 7]. After denervation, we observed decreases in the proportions of both type I and type II myonuclei (Figure 1F); concomitantly, the fractions of FAPs and macrophages increased, that of satellite cells remained unchanged, and endothelial cells and pericytes decreased.

To characterize cellular heterogeneity after denervation, we applied Monocle 3 to the entire dataset. This algorithm arranges each cell along a pseudotime based on transcriptional similarities, thus can determine the patterns of dynamic change experienced by different cell types. With our dataset, the most complex trajectory was generated in myonuclei, followed by FAPs and endothelial cells (Figure 1G, left panel). Globally, denervation elongated the trajectory of myonuclei, while the FAPs and endothelial trajectories were relatively shorter and contained fewer branches (Figure 1G, right panel). These results indicate that relative to other cell populations, denervation causes more significant changes in the heterogeneity and transcriptional profile of myofiber.

### snRNA-seq reveals heterogeneous transcriptional changes of type II myofiber in response to denervation

To delineate differences in the responses of different myofiber types to denervation, we isolated each subtype of myonuclei and used Monocle 3 to construct fiber-type-specific trajectories. We first constructed the trajectory of type II myonuclei, which contribute the most to myofiber composition in mouse Gas muscle (Figure 2A, left panel). In normal muscle, type II myonuclei formed a continuous trajectory extending from type IIa at one end to type IIb at the opposing end. Notably, type IIb myonuclei were further divided into two sub-groups, which we named type IIb1 and type IIb2 (Figure 2A, middle panel). Upon denervation, this continuum was disrupted by an obvious deviation of type IIb2 myonuclei, which formed a distinct cluster emerging at the end of the trajectory (Figure 2A, right panel). This response was accompanied by a remarkable increase in the proportion of type IIb2 myonuclei and decreases in other type II myonuclei (Figure 2B), implying that in type II myofibers, denervation promotes a conversion of myonuclei into type IIb2.

**Figure 2.**
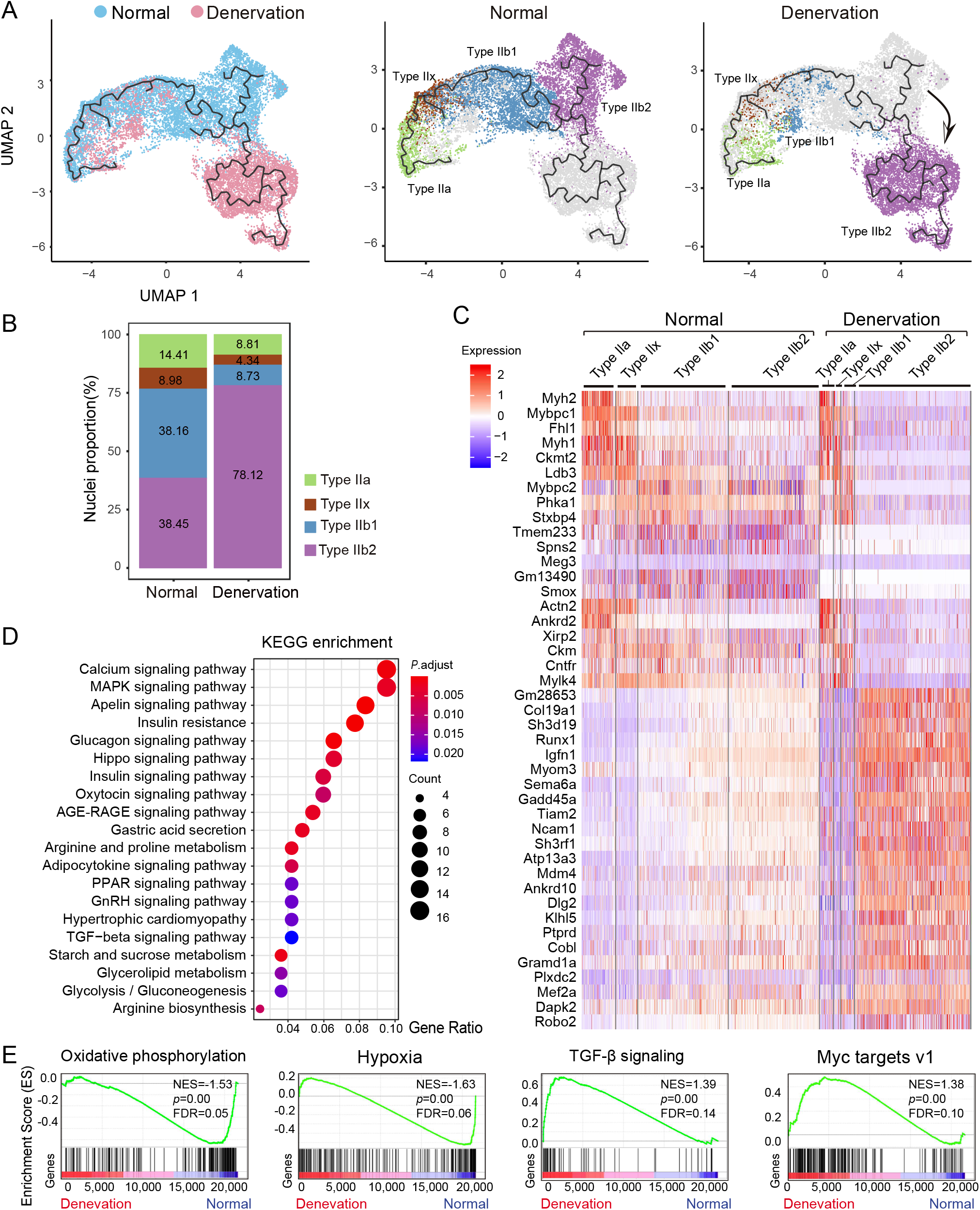
Trajectory illustrated transcriptional heterogeneity in type II myonuclei. (A) UMAP showing trajectory of type II myonuclei from normal (left panel, n=10561) and denervated muscles (right panel, n=6631). The myonuclei are colored according to conditions (left panel, blue: normal; pink: denervation) or their identified subtypes (middle and right panel). Type IIa: type IIa myonuclei; Type IIx: type IIx myonuclei; Type IIb1: type IIb1 myonuclei; Type IIb2: type IIb2 myonuclei (B) Proportion of different type II myonuclei subtypes in normal and denervated conditions. (C) Heatmap showing the signature genes of type II myonuclei subtypes. The color scale represents the relative expression level of gene in each nucleus.(D) Enriched KEGG pathways (p < 0.01) in denervated type IIb2 myonuclei. The color scale indicates the significance level of enrichment (adjust p value). Dot size represents counts of genes enriched in the pathway. KEGG: Kyoto Encyclopedia of Genes and Genomes. (E) GSEA plots showing enrichment score (ES) of the significant enriched hallmark gene sets in type IIb2 myonuclei. A positive value of ES indicates enriched in denervation condition, and a negative value indicates enriched in normal condition but downregulated in denervation. GSEA: gene set enrichment analysis; NES: normalized enrichment score; FDR: false discovery rate.

To characterize the denervated type IIb2 myonuclei, we identified their signature genes by comparing gene expression profiles with other type II myonuclei. We first surveyed the expression level of atrophic marker *Fbxo32* in type II nuclei upon denervation and found that it was significantly increased in all nuclei subtypes except type IIx nuclei, with the greatest expression level in type IIb2 nuclei (Figure S2A). We also found significant up-regulation of signature genes such as *Runx1, Gadd45a*, *Igfn1*, *Robo2*, *Dlg2*, and *Sh3d19* (Figure 2C). Runx1 and Gadd45a are reportedly involved in the pathogenesis of denervation-induced muscle atrophy [19, 20], while Igfn1 has been identified as essential for myoblast fusion and differentiation [21], but its role in denervated muscle remains unidentified. Likewise, the roles of many other signature genes in muscle atrophy have not been fully explored. For example, Robo2, a key component of the Robo–Slit signaling pathway with a profound role in embryotic myofiber formation [22], was highly up-regulated in denervated type IIb2 nuclei, implying a reactivation of embryonic myogenesis process in response to denervation. To locate the distribution of type IIb1 and IIb2 nuclei in muscle fibers, we performed immunofluorescence staining of Smox and Plxdc2, two signature genes of type IIb1 and IIb2, respectively. We observed a co-staining of these two proteins in the same type II fibers, suggesting that type IIb1 and IIb2 nuclei are present in the same fiber (Figure S2B).

We further performed pathway enrichment analysis on the differentially expressed genes (DEGs) identified in type IIb2 myonuclei using the Kyoto Encyclopedia of Genes and Genomes (KEGG) and found these DEGs to be primarily enriched in calcium signaling, MAPK, Hippo, and metabolic pathways (Figure 2D). Further gene set enrichment analysis (GSEA) revealed that hallmark gene sets involving oxidative phosphorylation and hypoxia were enriched in normal myonuclei but downregulated in denervated type IIb2 myonuclei. Conversely, hallmark sets with genes involving TGF-beta signaling and MYC targets were significantly enriched in denervated type IIb2 myonuclei (Figure 2E, Figure S2C).

Lastly, we surveyed the activity of gene regulatory networks (GRNs) in type II myonuclei using single-cell regulatory network inference and clustering (SCENIC) [16]. This approach maps and scores the activity of regulons, which consist of transcription factors (TFs) and their regulated target genes, to determine the underlying GRNs in a given cell type. We identified 147 active regulons in normal and denervated type II myofibers (Figure S3A). Figure 3A presents the regulons whose activities were dramatically changed by denervation in type II myonuclei, alongside a list of master regulators that may mediate the transcriptomic and phenotypic changes of denervated type II myofibers. Using the regulon activity score of each nucleus, all myonuclei could be divided into the expected subtypes (Figure 3B), consistent with the clustering results obtained using cell identities and trajectory analysis (Figure 2A). The results from applying t-distributed stochastic neighbor on the binary regulon activity matrix also supported that denervation dramatically changes regulon activity in type II myonuclei. For example, the Hlf and Maf regulons were active in all normal type II myonuclei but were specifically suppressed in denervated type IIb2 nuclei; in contrast, the Elk4, Bhlhe40, Meis1, and Zfp369 regulons were predominantly enhanced in denervated type IIb nuclei, especially in the typeIIb2 subtype (Figure 3C). Figure S3B presents other regulons whose activities were enhanced by denervation, including Runx1, Mef2a, Myog, Myod1, and Myf6. These results indicate that distinctive gene regulatory networks were activated in type IIb1/2 myonuclei, driving the fiber type-specific transcriptional changes observed after denervation.

**Figure 3.**
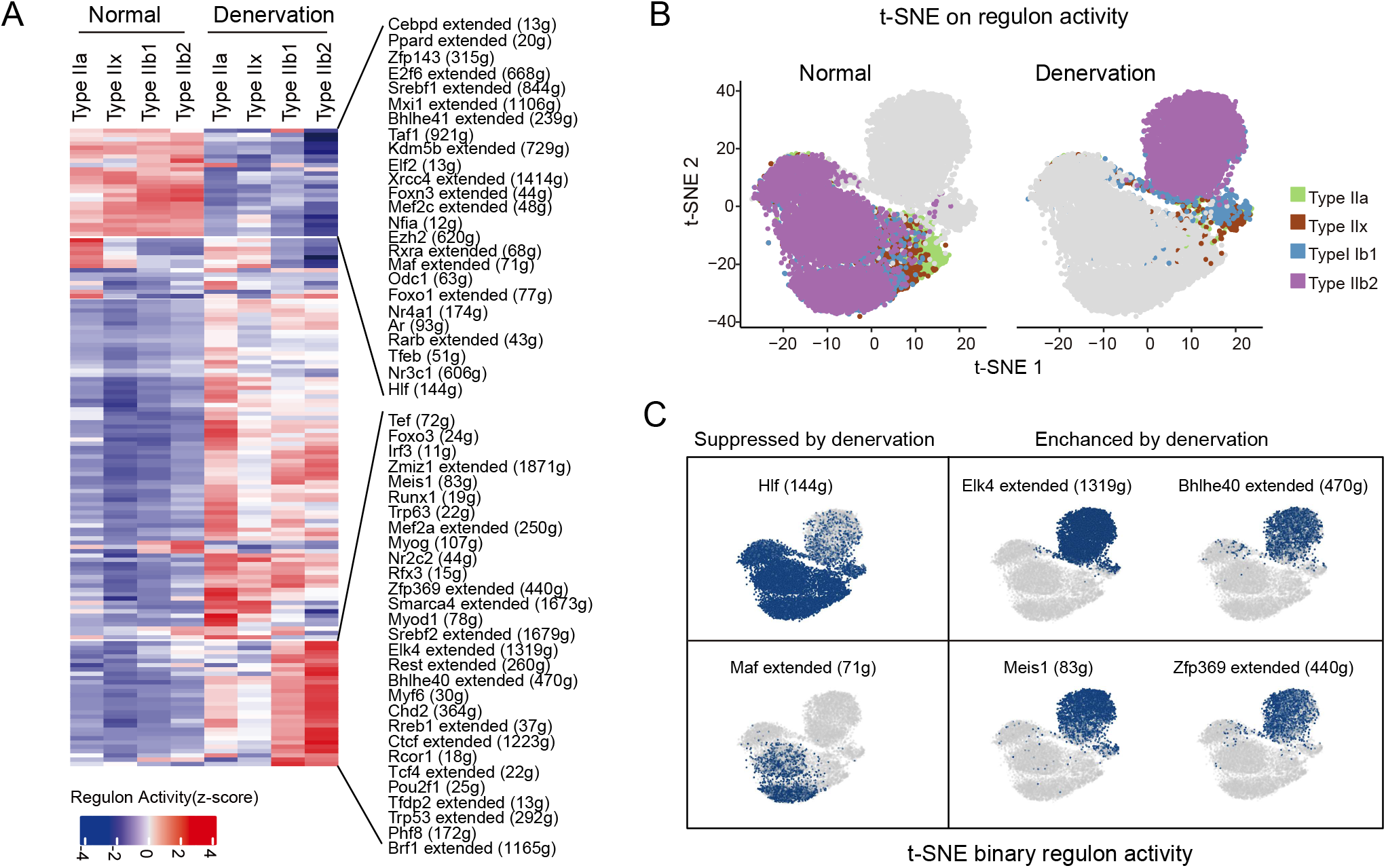
Gene regulatory network (GRN) of type IIb2 myonuclei in normal and denervation. (A) Heatmap showing regulons with significant changes between normal and denervated type IIb2 myonuclei. The color scale represents the activities of regulons. The top changed regulons and the counts of downstream target genes in the regulons are provided on the left. (B) t-distributed stochastic neighbor embedding (t-SNE) on binary regulon activity matrix of normal (left panel) and denervated (right panel) type II myonuclei. Each nucleus is colored according to corresponding subtype. (C) t-SNE on the binary regulon activity matrix showing denervation suppresses (left panel) or enhances (right panel) the activities of select regulons in myonuclei.

### Transcriptomic and heterogeneous alternations in denervated type I myofiber

To dissect the transcriptomic changes of type I fiber in response to denervation, we reconstructed the trajectory of type I myonuclei. Unlike type II myonuclei, the trajectory of type I myonuclei under normal conditions had no nodes or branches, indicating lower heterogeneity within type I myonuclei. In response to denervation, the affected type I nuclei “drifted away” from their normal counterparts and formed a distinct cluster (Figure 4A), indicating that denervation dramatically changed their transcription profiles and supporting the notion that type I myofibers are profoundly affected by denervation [23]. We then analyzed DEGs between these two distinct clusters, identifying 379 genes that were significantly changed. Among these, the top up-regulated genes were *Dlg2*, *Col25a1*, *Igfn1*, *Col23a1*, *Arpp21;* while the top down-regulated genes were *Rp1*, *Hs3st5*, *Rnf150*, *Mylk4*, and *Lrrfip1* (Figure 4B). Other relevant DEGs are shown in Figure S4A. Notably, those significantly up-regulated genes included oxidative stress-responsive genes such as *Car3* and *Sfpq*. Using KEGG analysis, we found the DEGs of denervated type I myonuclei were related to pathways involving protein degradation, autophagy, AMPK, mTOR, and insulin signaling (Figure 4C). GSEA analysis revealed that gene sets for fatty acid metabolism, oxidative phosphorylation (OXPHOS), PI3K/mTOR pathway, and adipogenesis were significantly enriched in normal but downregulated in denervated type I myonuclei (Figure 4D and Figure S4B). Furthermore, SCENIC analysis revealed the dominant gene regulatory networks in normal type I fibers include Mta3, Nr1h3, Rxrg, and Yy1 regulons. In response to denervation, the activities of these regulons decreased while those of the C/EBPs, Sox13, Nr2c1, Mga, and Stat6 regulons were dramatically increased (Figure 4E). In particular, CEBP-β, δ, γ, and ζ regulons were remarkably activated in denervated type I nuclei. A prior report supports that C/EBPβ promotes the expression of atrogenes in cancer cachexia [24]; together with our results, these findings suggest that GRNs driven by C/EBPs are essential for the pathogenesis of muscle atrophy. Another activated regulon is Sox13, however the function of Sox13 in skeletal muscle has not been explored. Another TF of interest is Mta3, which reportedly regulates the epithelial-to-mesenchymal transition [25] and stimulates the expression of HIF1 under hypoxia conditions [26]. Yet little is known about its role in skeletal muscle biology and the pathogenesis of muscle atrophy.

**Figure 4.**
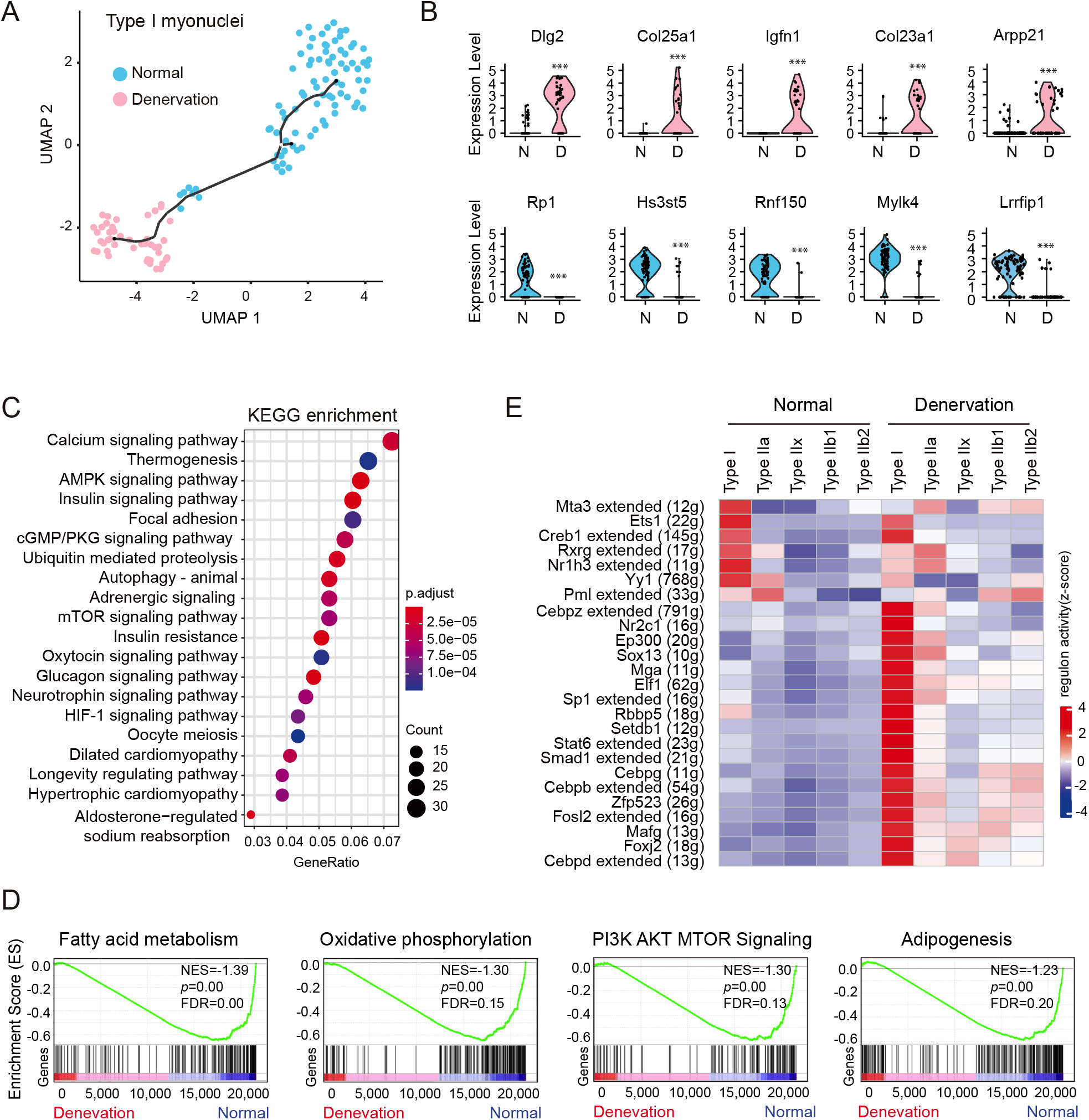
Transcriptional responses of type I myonuclei to denervation. (A) UMAP showing the pseudotime trajectory of type I myonuclei from normal (blue, n=88) and denervated (pink, n=44) muscles. (B) Violin plot showing the expression of select differentially expressed genes (DEGs) between normal (blue) and denervated (pink) type I myonuclei. N: normal; D: denervation. Wilcoxon Rank Sum test: min.pct = 0.25, logfc.threshold = 0.25. Significance level: *** p < 0.001. (C) GSEA plots showing enrichment score (ES) of the significant enriched hallmark gene sets in type I myonuclei. NES: normalized enrichment score; FDR: false discovery rate. (D) Enriched KEGG pathways (p < 0.01) of DEGs in denervated type I myonuclei. The color scale indicates the significance level of enrichment (adjust p value). Dot size represents counts of genes enriched in the pathway (E) Heatmap showing top changed regulons between normal and denervated type I myonuclei. The color scale represents the activities of regulons. The number in parentheses represents the count of target genes in corresponding regulons.

### Metabolic plasticity and heterogeneity of myonuclei after denervation

Different muscle fiber types exhibit unique metabolic properties; accordingly, we characterized the metabolic heterogeneity of myofibers and their respective adaptive responses to denervation at single nucleus resolution. Specifically, we compared the normal and denervated metabolic gene expression profiles of each subtype using the Single-Cell-Metabolic-Landscape pipeline [17]. This approach calculates a metabolic pathway activity score according to the relative gene expression averaged over all genes in that pathway. We first performed clustering of the 1401 detected metabolic genes in type I/II myonuclei (Figure 5A, 5B) and the result was similar to the classification using the complete set of genes (Figure 1A), indicating metabolic features weighs heavily on the phenotypic changes of fiber types. We next quantified the overall metabolic pathway activities of normal and denervated cells based on myonuclear type. For normal cells, we observed a gradually decreasing trend in metabolic activity from oxidative type I myonuclei to glycolytic type IIb2 myonuclei. Upon denervation, we found that all myonuclei exhibited significant down-regulation of metabolic activity, reflecting the reduced metabolic demands of denervated myofibers (Figure 5C). The affected metabolic pathways mainly concerned lipid, carbohydrate, amino acid, and nucleotide metabolism, along with the citrate cycle and oxidative phosphorylation (Figure 5D). These findings are in line with reports that denervation induces massive metabolic adaptation in muscles.

**Figure 5.**
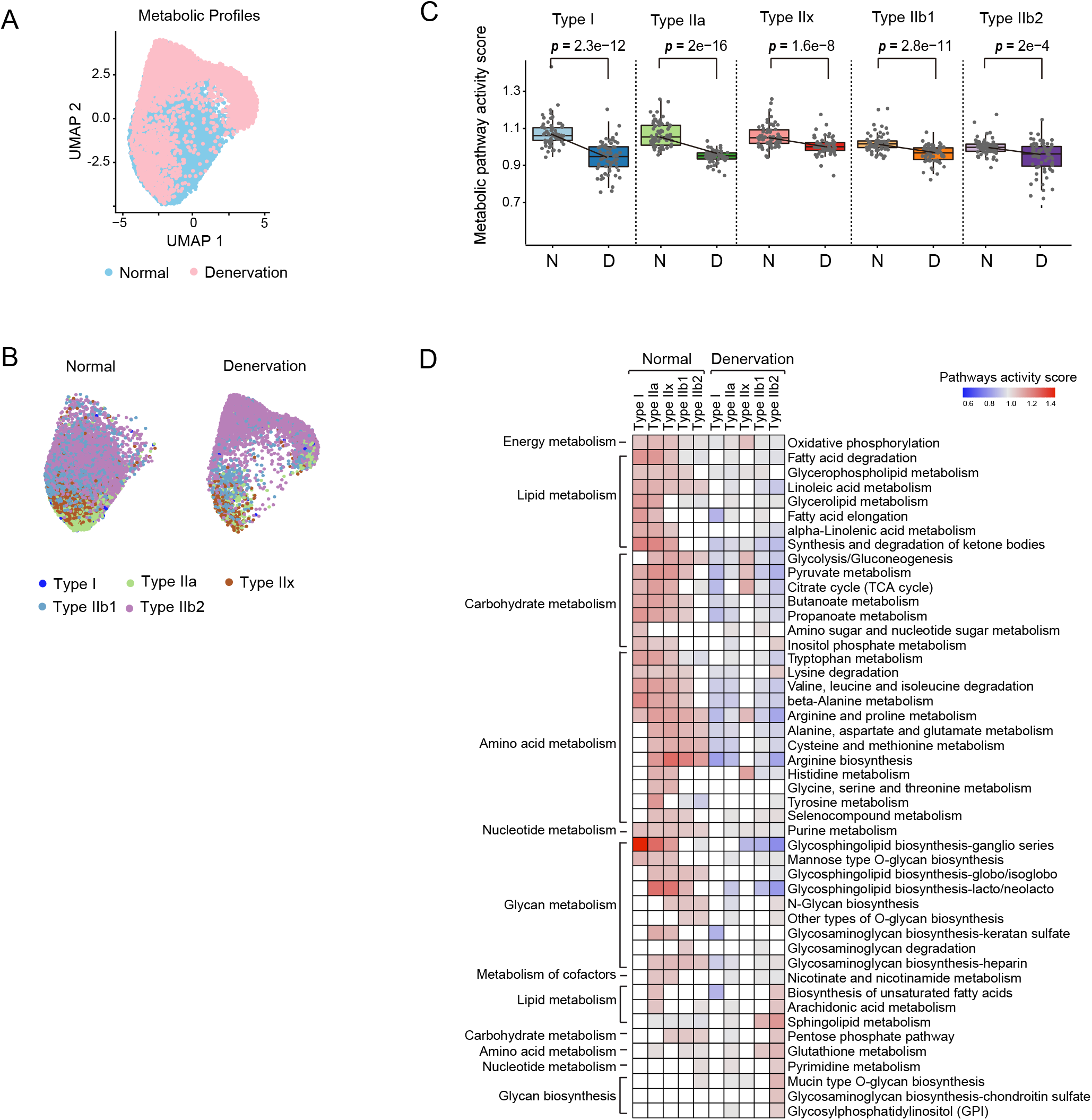
Metabolic landscape of myonuclei from normal and denervated muscles. (A) Clustering on metabolic gene expression profiles of myonuclei enables identification of normal and denervation cell states. The dots are colored according to different conditions (normal, blue; denervation, pink). (B) UMAP of metabolic profiles of different myonuclei subtypes in normal (right panel) and denervation (left panel). The dots are colored according to corresponding subtypes. (C) Boxplot showing the metabolic activities of different myonuclei subtypes between normal and denervation. Each dot represents the activity score of an individual metabolic pathway. Comparisons between conditions are performed using Wilcoxon rank-sum test. N: normal; D: denervation. (D) The activity score of top metabolic pathways detected in myonuclei subtypes. The blank indicates that the score of pathway activity is not significant in relevant subtype (random permutation test, p > 0.01). The metabolic pathways are arranged according to categories.

Intriguingly, the magnitude of decline in metabolic gene expression induced by denervation decreased from oxidative to glycolytic myonuclei: that is, the largest decrease was observed in type I nuclei and the smallest in type IIb2 nuclei. This suggests that type I myofibers are more vulnerable to denervation; however, we also observed denervated type IIb2 myonuclei to feature striking up-regulation of metabolic pathways involving unsaturated fatty acid, sphingolipid, pyrimidine, and glutathione metabolism (Figure 5D). Sphingolipid and unsaturated fatty acid metabolism are closely linked to muscle biology and pathophysiology such as muscle protein loss and insulin resistance [27]; meanwhile, glutathione and pyrimidine derivatives are important antioxidants, and increased activity of associated pathways implies a compensatory response to oxidative stress. We also observed an increase in activity for glycan biosynthesis, which is important for the modulation of cell communication and signaling transduction; however, the role of this metabolic component during muscle denervation needs further exploration.

### Heterogeneity and phenotype of FAPs in response to denervation

The second complex trajectory was formed by FAPs. In normal, this trajectory was restricted to a concentrated area and segregated into three subclusters (C1-3) (Figure 6A, upper panel). Upon denervation, those subclusters went through remarkable expansions to form three branches emanating from the main trajectory (Figure 6A, lower panel), indicating that FAPs undergo substantial transcriptome changes. To characterize these changes, we isolated FAPs from each branch and compared their transcriptomes with those of the corresponding normal subcluster. The top 20 DEGs in each subcluster are listed in Figure S5A. In addition, we surveyed the expression patterns of known FAPs markers in each cluster (Figure S5B), and determined them to be consistent with the results of a published study [5]. We then performed GSEA and compared the hallmark profiles of each subcluster under normal and denervation conditions. In normal condition, all three subclusters showed significantly enriched hallmarks related to myogenesis, hypoxia, and metabolism, which were downregulated under denervation conditions. After denervation, these subclusters were found to be enriched for different gene sets (Figure 6B). The denervated C1 subcluster was enriched for hallmarks of apoptosis, coagulation, and the P53 pathway, suggesting that this subcluster might represent an apoptotic phenotype [28]. Meanwhile, the C2 subcluster was enriched for hallmarks related to epithelial-mesenchymal transition (EMT), angiogenesis, and TGF-β signaling, and the C3 subcluster for adipogenesis, MYC targets V1, and Wnt/beta-catenin signaling. These results imply that denervation activates C2 and C3 FAPs to acquire pro-fibrotic and pro-adipogenic features, respectively (Figure 6A, lower panel).

**Figure 6.**
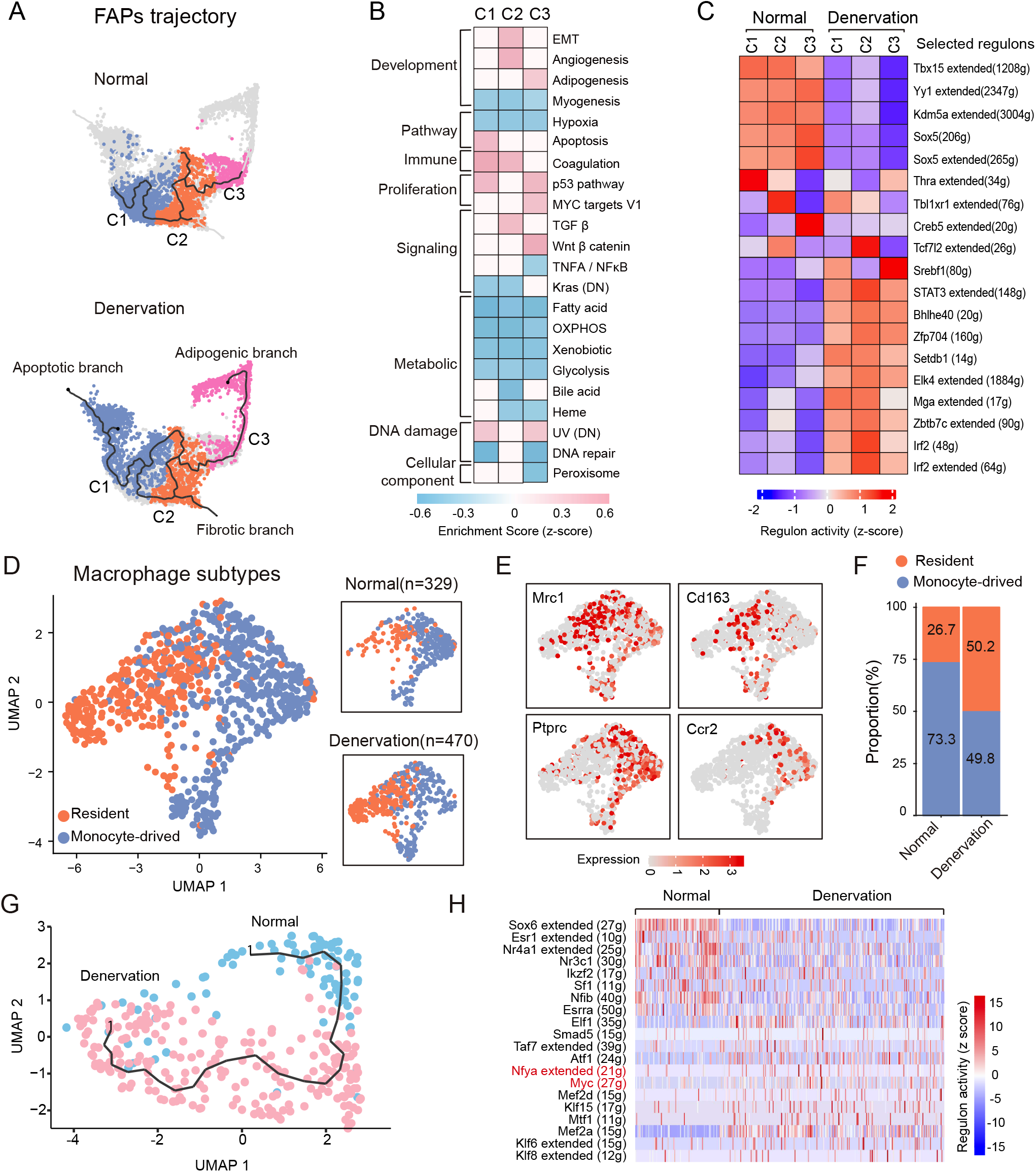
Transcriptional responses to denervation in Fibro-adipogenic progenitors (FAPs) and macrophages. (A) UMAP showing a branching trajectory of FAPs from normal muscles (upper panel, n=2825) and denervated muscles (lower panel, n=3634). The nuclei mapped on the path are colored and labeled according to their identified subtypes as C1, C2, and C3. These subclusters are annotated as apoptotic branch, fibrotic branch and adipogenic branch according to GSEA analysis. (B) Heatmap showing GSEA enrichment scores of the 3 FAPs subtypes in denervated muscle compared with corresponding normal subtypes. The category annotations of hallmark gene sets are based on information from GSEA database. A positive enrichment score (ES) indicates the gene set is significantly enriched in denervated FAPs, and a negative ES indicates that gene set is significantly enriched in normal FAPs. The blank indicates there is no significant enrichment of the gene set (p >0.05, FDR >25%). (C) The most active regulons in 3 subtypes of FAPs from normal and denervated muscles. The color scale represents the normalized scores of regulon activity: red denotes high level of activity while blue indicates low level of activity. The number in parentheses represents the count of downstream target genes in corresponding regulons. (D) Left panel: UMAP showing 2 subclusters of macrophages colored with identified phenotypes. Resident: skeletal muscle resident macrophages (orange); monocyte-derived: monocyte-derived macrophages (blue). Right panel: Visualization of the 2 macrophages subtypes on UMAP split by conditions (normal and denervation). The number in brackets denotes the total numbers of macrophage from normal and denervated muscles. (E) Visualization of the expression and distribution of established macrophage identities in 2 subtypes of macrophage. (F) Changes in proportions of macrophages subtypes in normal and denervated muscles. Different colors represent the corresponding subtypes. (G) Trajectory of resident macrophages from normal and denervated muscles. Each dot represents a nucleus and is colored according to conditions (blue, normal; pink, denervation). (H) The activity scores of top20 regulons detected in normal and denervated resident macrophages.

We then inferred the GRNs underlying the transcriptomic and phenotypic changes of FAPs and found 297 active regulons in normal and denervation states (Figure S6A). In normal, all subtypes of FAPs were found to be governed by some common regulons including Kdm5a, Tbx15, Yy1, and Sox5; however, unique patterns of regulon activity were also observed in different subclusters, such as higher activities of Thra (C1), Tbl1xr1 (C2), and Creb5 (C3) regulons (Figure 6C). Upon denervation, most active regulons in normal FAPs were suppressed, with concomitant adaptive activation of a diverse range of other regulons; in particular, over 120 regulons were strongly activated in the C3 subcluster (Figure S6A). It was notable that some specific regulons potentially drive the phenotypic changes of FAPs, such as theTcf7l2 and Srebf1 regulons activated in C2 and C3 FAPs after denervation. Tcf7l2 plays a major role in the Wnt signaling pathway, which regulates fibrosis [29], while Srebf1 is a key regulator of adipogenesis [30]. We also found that C2 FAPs exhibited higher activity of the Stat3 regulon, which was reported to promote myofiber atrophy and fibrosis during denervation [9]. Finally, we highlight several novel regulons whose activities were robustly increased in FAPs after denervation (Figure S6A), the roles of these TFs during muscle atrophy merit further investigation.

### Alteration of macrophages in denervated muscles

In both normal and denervated muscles, macrophages (MPs) were the predominant inflammatory cells we captured (Figure 1C). These cells were further divided into two distinct subclusters, mainly representing skeletal muscle-resident MPs (*Mrc1*- and *Cd163*-positive) and monocyte-derived MPs (*Ptprc*- and *Ccr2*-positive) (Figure 6D, 6E). Detailed signatures gene for both groups are provided in Figure S7A. Notably, in denervated muscles, we observed an increase in total numbers of macrophages and a remarkably higher proportion of resident MPs (Figure 6D, 6F). Thus, we focused further analysis on resident MPs. In trajectory analysis, we found resident MPs generated a continuous trajectory without branches, with its two clusters corresponding to normal and denervated resident MPs (Figure 6G). Analysis of DEGs revealed that in normal muscles, resident MPs expressed higher levels of *Tmem233*, *Ano5*, *Mybpc*, and *Capn3*. In the context of denervated muscles, those genes were significantly down-regulated, while genes relating to fibrosis or immune responses, such as *Col19a1*, *Ncam1*, and *Enpp3*, were up-regulated (Figure S7B). GSEA analysis revealed gene sets involving immune response, chemokine receptor binding to be significantly enriched in denervated MPs (Figure S7C). Finally, we used SCENIC to identify the regulators responsible for the transcriptional changes in resident MPs between normal and denervation conditions. We identified several key regulons for normal MPs, including Nr4a1, Nfib, and Sox6, and for denervated MPs, such as Mef2a, Smad5, Myc, and Nfya (Figure 6H). Since Myc and Nfya are reportedly activated in self-renewing macrophages [31], these results imply that denervation may enhance the self-renewal capacity of resident macrophages.

### The transcriptome and heterogeneity of muscle stem cell (MuSCs) within denervated muscle

To precisely depict the transcriptomic changes between normal and denervated MuSCs nuclei, we projected these nuclei onto a UMAP to visualize their distribution and relationship. As described by Figure 7A, these nuclei formed two separated clusters; each cluster contains nuclei from both normal and denervated conditions. We then performed DEG analysis and found the expression of Sorbs2, Ppm1l, Pygm, Pdk4, Amd1 were significantly downregulated in denervation (Figure 7B). We also interrogated the expression of satellite cells markers in both clusters. The markers for quiescent satellite cells (Pax7, Cav1, Spry1, Btg2) were detected in normal; the markers for activated satellite cells (Myod1, Myog, Myf5, Desmin, Numb) were detected in denervated MuSCs. Genes that regulate satellite cell self-renewal (Myf5, Par-3, Six1) were also observed in denervated MuSCs. Notably, the transcripts of cell cycle genes (such as Ccna2, Ccnb1, Ccnb2, Ccne1) were undetectable in MuSCs from both conditions (Figure 7C). We further characterized the biological process of DEGs in denervated MuSCs and found that these DEGs mainly enriched in PI3K-AKT and MAPK signaling pathway, protein digestion and absorption, regulation of actin cytoskeleton, focal adhesion and axon guidance (Figure 7D). Moreover, GSEA analysis revealed significant downregulation of muscle filament sliding, nucleotide metabolism and glycolysis in denervated MuSCs (Figure 7E). Instead, the enrichment of mesenchymal epithelial transition gene set was detected in denervated condition. Regulon activity analysis indicated that Ar, Nr3c1, Zbtb20 and Mef2c were the major regulons of normal MuSCs, whereas their activities were largely repressed in denervated MuSCs. On the other hand, Runx1, Ets1 and Hcfc1 regulons were enhanced upon denervation (Figure 7F and 7G). Based on previous studies, Mef2c can induce satellite cell proliferation and differentiation [32], and Zbtb20 were described to be induced in muscle stem cells and induce myogenesis [33]. Taken together, although MuSCs show signs of activation in response to denervation, their myogenesis ability was impaired in denervated muscle.

**Figure 7.**
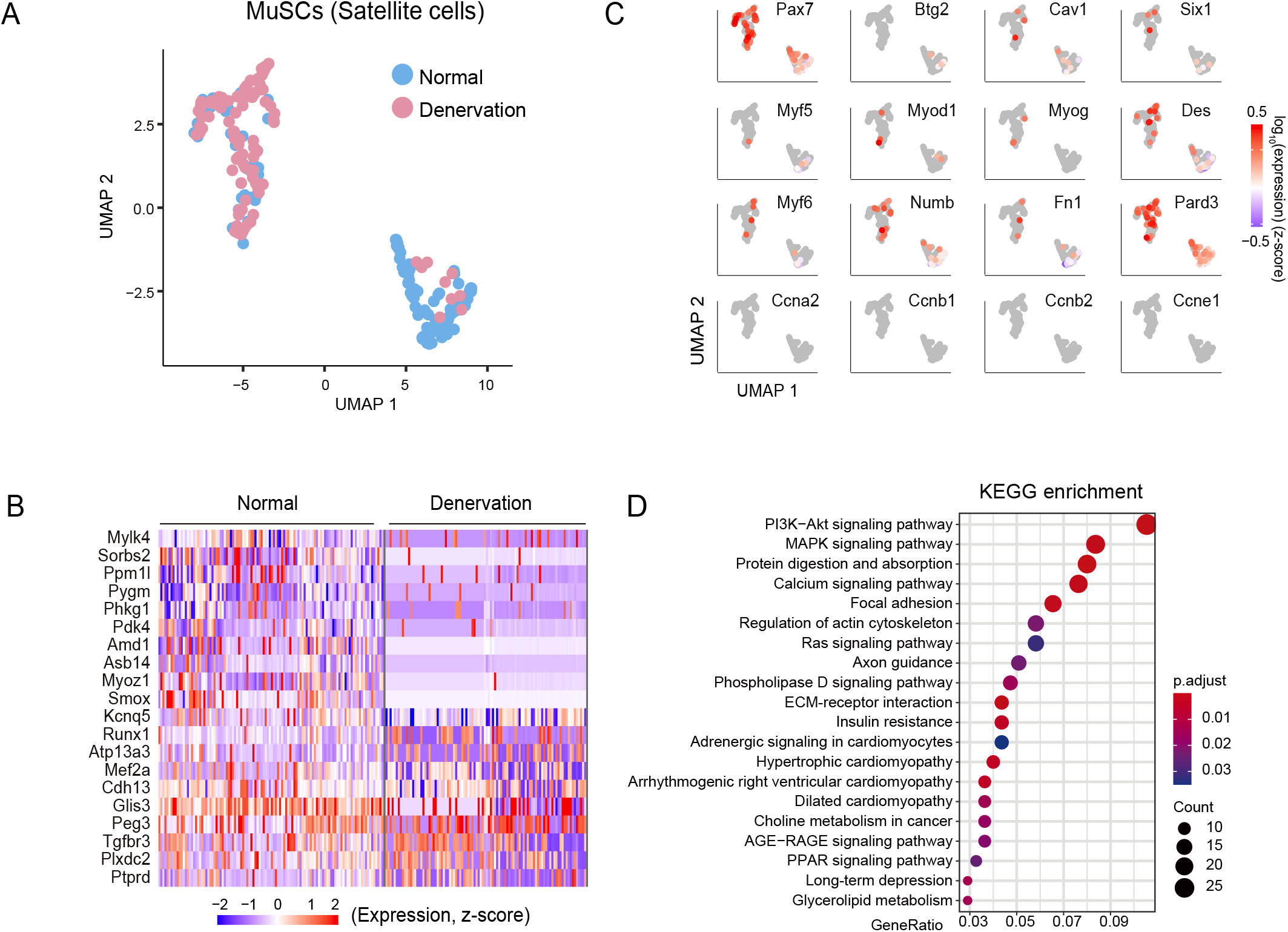
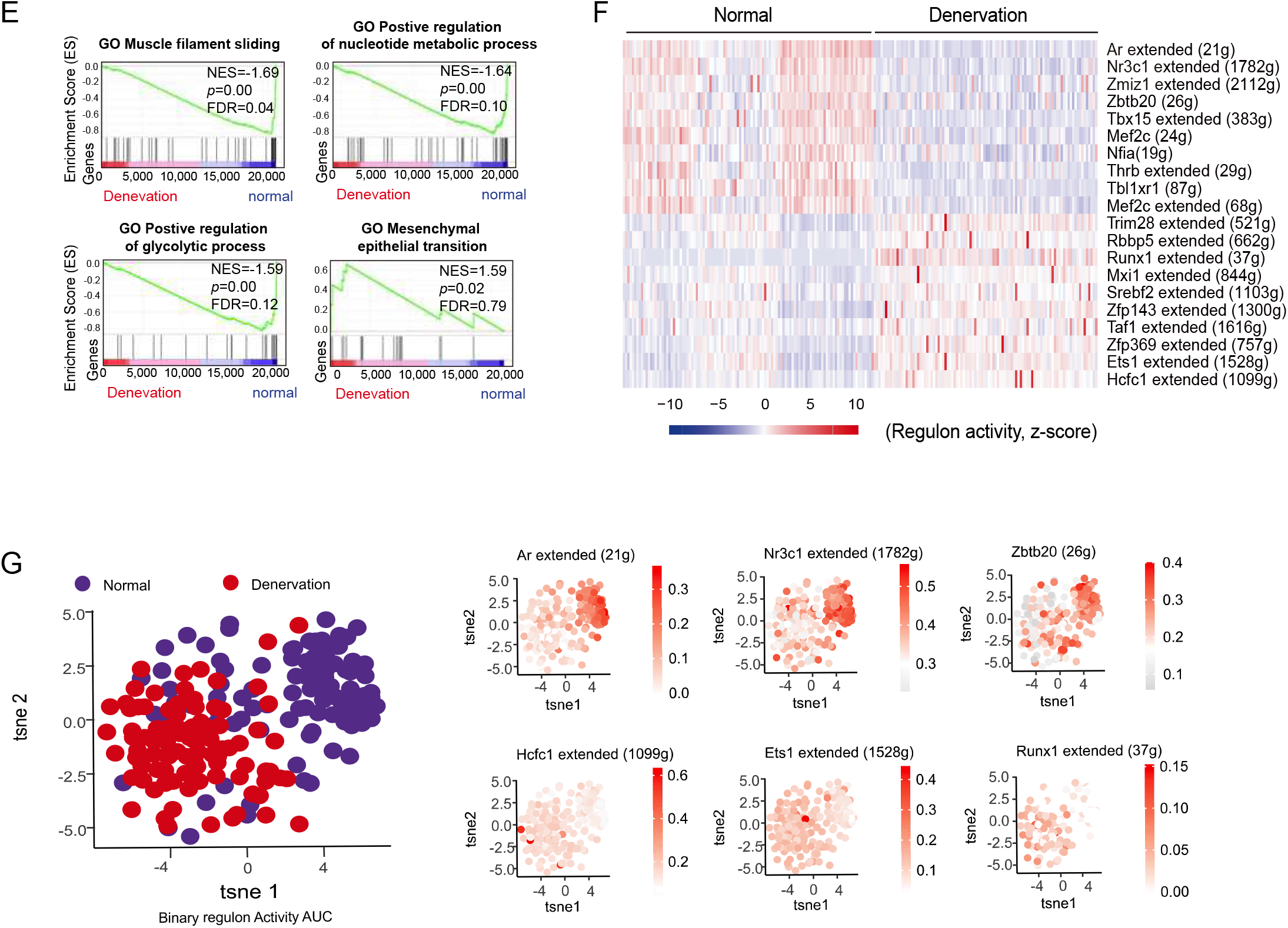
Transcriptomic responses of muscle stem cells (MuSCs) to denervation. (A) UMAP showing the trajectory of MuSCs from normal (blue, n=110) and denervated (pink, n=98) muscles. (B) Heatmap displaying the top 20 DEGs between normal and denervated MuSCs. The color scale represents the relative level of gene expression: dark blue indicates low level of and red indicates high levels of expression. (C) Changes in expression of select genes in MuSCs along the pseudotime trajectory. Each point indicates a nucleus. The color intensity represents log2 expression value of the expressed gene; darker color indicates higher expression level. (D) Significantly enriched KEGG pathways in MuSCs from denervated muscles (p < 0.05). The color scale indicates the significance level of enrichment (adjust p value). Dot size represents counts of genes enriched in the pathway. (E) GSEA plots showing the significantly enriched GO gene sets in MuSCs. GO, Gene Ontology; NES: normalized enrichment score; FDR: false discovery rate. (F) Heatmap showing top changed regulons between normal and denervated MuSCs. The color scale represents the normalized scores of regulon activity: dark blue indicates low level of activity and red indicates high level of activity. The number in parentheses represents the count of downstream target genes in corresponding regulons. (G) Left panel: t-SNE on binary regulon activity matrix identifying two closely related clusters, representing normal (purple) and denervated (red) MuSCs, respectively. Right panel: t-SNE showing active regulons in MuSCs form normal and denervated muscles. Each dot represents a nucleus with high regulon activity.

### Transcriptional features of endothelial cells and pericytes in denervated muscles

During muscle atrophy, vascular endothelial cells (ECs) exhibit a massive adaptive response [34]. To explore the impact of denervation on these cells, we first sub-clustered them, identifying four subtypes. In addition to capillary ECs and lymphatic ECs, we found two subclusters expressing both a capillary marker and either an arterial or a venous marker, which may respectively represent arteriole and venule ECs (Figure 8A). Cell identities for these classifications are provided in Figure S8A. Under normal conditions, capillary ECs were the dominant subtype; however, after denervation, the proportion of capillary ECs decreased greatly (76.7% vs 26.2%), while the relative proportions of arteriole and venule ECs increased, and the proportion of lymphatic ECs remained almost unchanged (Figure 8B). Furthermore, we found the EC subtypes to exhibit heterogeneous responses to denervation. As delineated by the cellular trajectory, denervation stimulated the vascular ECs to undergo substantial transcriptomic changes, forming two distinct groups interposed by lymphatic ECs (Figure 8C). These two separated groups mainly represented the normal and denervated capillary ECs. Due to capillary ECs exhibiting significant changes in both abundance and transcriptomic profile following denervation, we focused on this subtype and characterized DEGs in terms of their associated biological processes. Pathway analysis revealed the DEGs to be enriched in several pathways, including calcium signaling, the MAPK pathway, ubiquitin mediated proteolysis, and the glucagon and insulin related pathway (Figure 8D). GSEA analysis showed that most of the gene sets enriched in normal ECs were down-regulated upon denervation, including hypoxia, oxidative phosphorylation, and metabolism-related hallmarks (Figure 8E).

**Figure 8.**
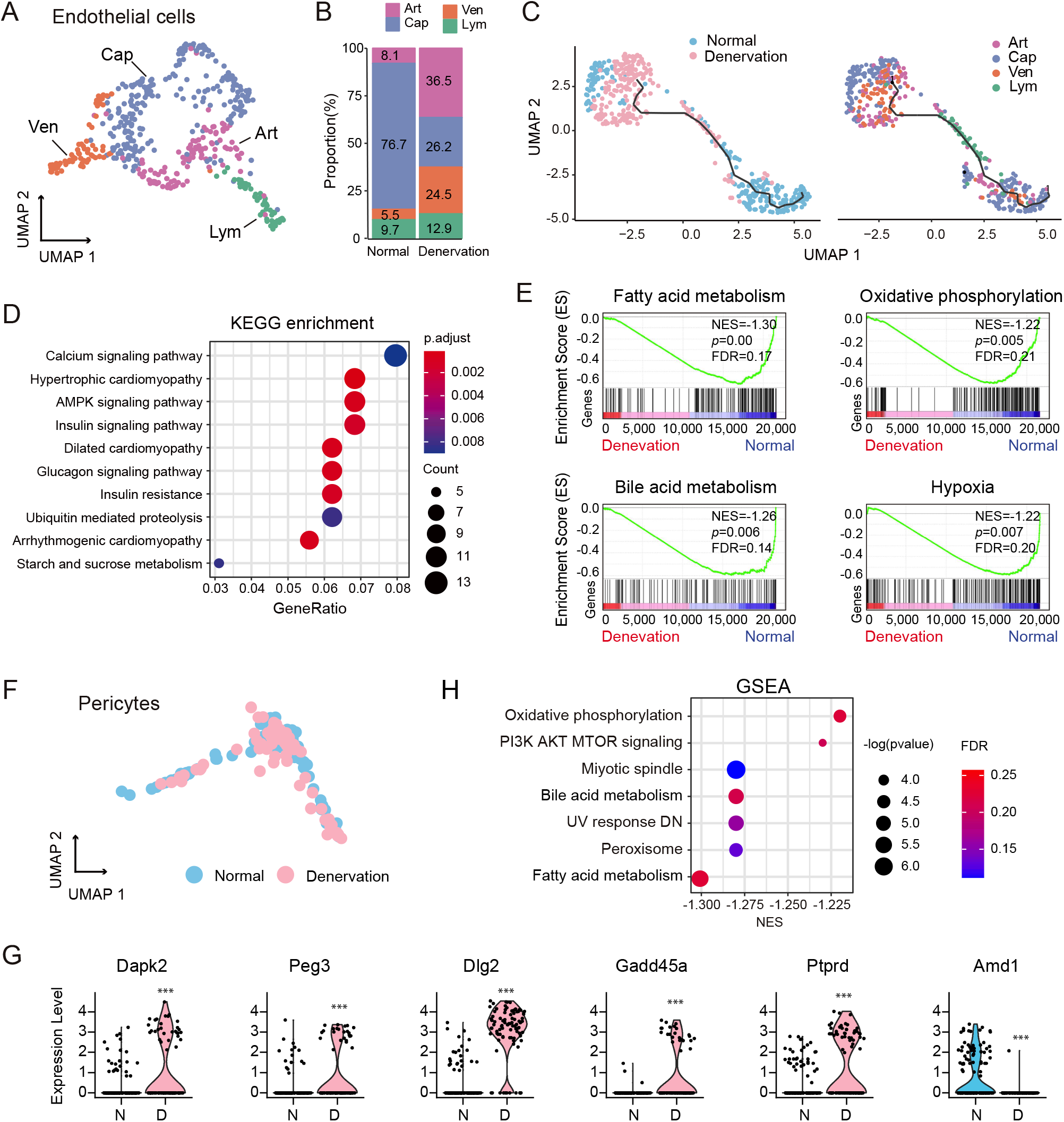
Transcriptional heterogeneity in endothelial cells (ECs) and pericytes in response to denervation. (A) UMAP visualizing 4 subclusters of ECs colored according to their identified subtypes. Art: arteriole endothelial cells; Cap: capillary endothelial cells; Ven: venule endothelial cells; Lym: lymphatic endothelial cells. (B) Changes in proportions of ECs subtypes from normal and denervated muscles. Different color codes represent the corresponding subtypes described as (A). (C) Trajectory of ECs from normal and denervated muscles. Left panel: The nuclei mapped on the path are colored according to conditions (normal, blue; denervation, pink). Right panel: The nuclei mapped on the path are colored by identified subtypes. (D) Significant enriched KEGG pathways (p < 0.01) of DEGs between normal and denervated capillary ECs. The color scale indicates the significance level of enrichment (adjust p value). Dot size represents counts of genes enriched in the pathway. (E) GSEA plots showing enrichment score (ES) of the significant enriched hallmark gene sets in capillary ECs. NES: normalized enrichment score; FDR: false discovery rate. (F) The trajectory of pericytes from normal (blue) and denervated (pink) muscles. (G) Violin plot displaying the difference in expression of select DEGs of between normal (blue) and denervated (pink) pericytes. N: normal; D: denervation. Wilcoxon Rank Sum test: min.pct = 0.25, logfc.threshold = 0.25. Significance level: *** p < 0.001. (H) The significant enriched hallmark gene sets identified by GSAE analysis in pericytes. Dot size indicates the significance level of enrichment (-log (p value)). The color scale represents FDR value

Regarding pericytes, UMAP showed that these cells exhibited only subtle transcriptional differences between normal and denervated conditions (Figure 8F). DEG analysis revealed denervated pericytes to express higher levels of apoptosis-associated genes (*Dapk2* and *Peg3*) and stress response genes (*Gadd45a*, *Dlg2*, *Ptprd*, and *Amd1*) (Figure 8G). GSEA analysis also suggested that PI3K signaling, oxidative phosphorylation, and metabolic hallmarks were enriched in normal pericytes but dramatically downregulated in denervated pericytes (Figure 8H).

### Communication between myofibers and other resident cells in response to denervation

To infer the responses of intercellular communication to denervation at single-nucleus level, we applied a ligand-receptor (L-R) interaction model [3] to calculate scores for L-R pairs based on the expression levels of ligands and receptors. We first referred to a prior L-R interaction database [18] and observed extensive interactions between myofibers and resident cells (Figure S9A). In total, we predicted 592 and 21 potential L-R interaction pairs in normal and denervated conditions, respectively. Since the catabolic signaling induced by denervation is initiated in myofibers, we prioritized communications dominated by myofibers; specifically, we focused on ligands differentially expressed in myofibers and their corresponding receptors expressed in resident cells. In the normal condition, type I myofibers showed higher expression of the ligands *Mapt*, *Fgf1*, *Angpt1*, *Ptprm*, *Ncam1*, and *Jam3*, while type II myofibers had higher levels of *Fgf13*, *Cadm1*, *Egf*, and *Vegfa* (Figure 9A, left panel). Upon denervation, we found a dramatic dialing-down of intercellular L-R interactions, with satellite cells becoming the dominant cell type receiving signals from myofibers (Figure 9A, right panel and Figure S9B). The switchover included the decline of several ligands (such as *Fgf1* in type I fibers and *Vegfa* in type II fibers) and up-regulation of particular ligands in type II fibers (such as *Fgf13*, *Mapt*, and *Ncam1*) (Figure 9B). We examined the expression levels of these 3 genes in different subgroups of type II nuclei from denervated muscles and found that their levels in type IIb2 nuclei were significantly higher than other type II nuclei (Figure 9C).

**Figure 9.**
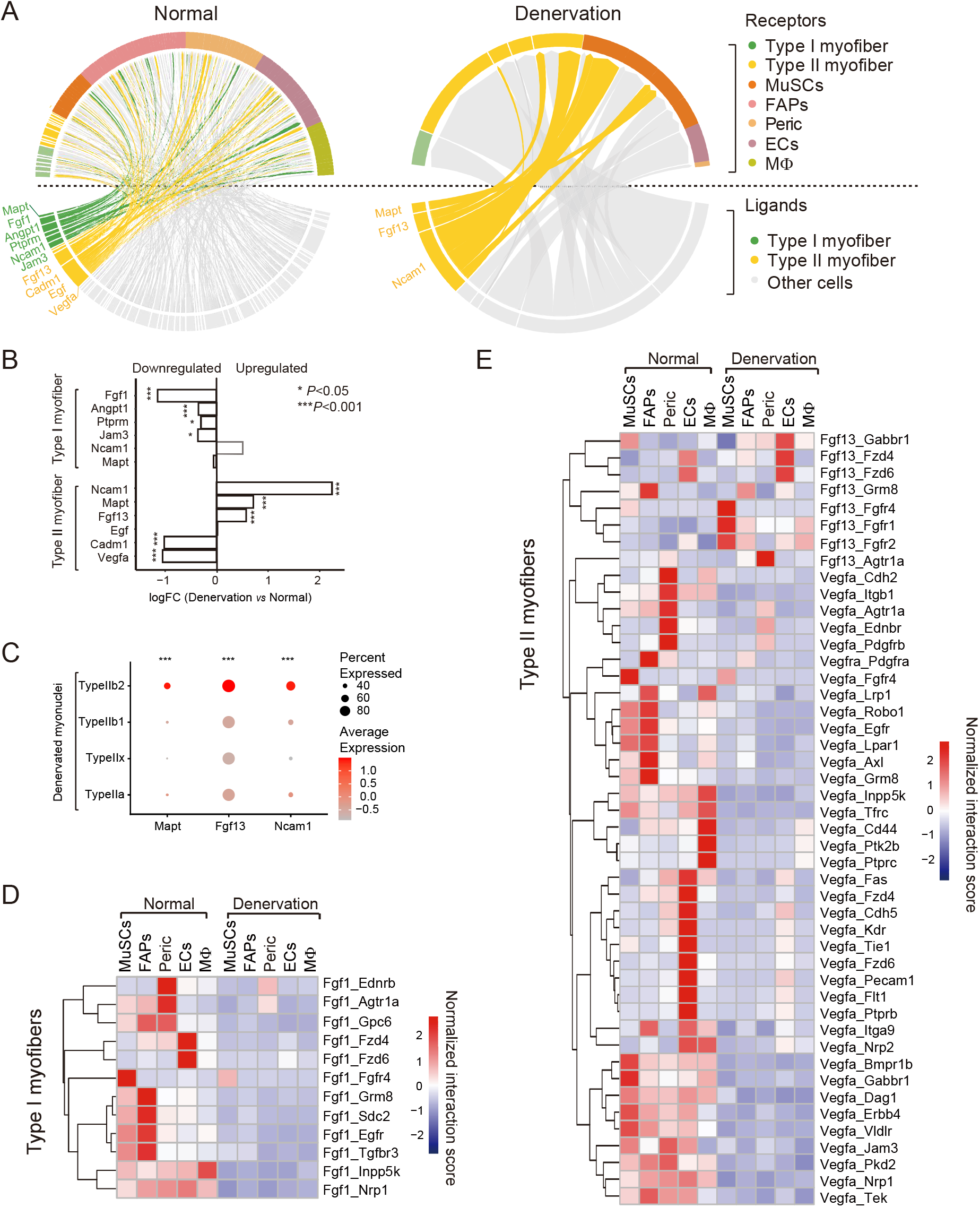
Ligand-receptor interactions between myofibers and other resident cells in response to denervation. (A) Chrod plot displaying intercellular ligand-receptor (L-R) interactions in normal (left panel) and denervated (right panel) muscles. Ligands differentially expressed by myofibers are listed below the dash line, and receptors differentially expressed by receiver cells (including myofibers and other types of resident cells) are listed above the dash line. The interactions between myofibers (L) and other resident cells (R) are highlighted (green and yellow). Other interactions are presented in gray. (B) Changes in expression level of ligands expressed by type I and type II myofibers between normal and denervated condition. Wilcoxon rank-sum test, significance level: *p <0.05, ***p <0.001. (C) Dot plot showing the expression levels of ligands i.e., Fgf13, Mapt, and Ncam1, in subgroups of type II nuclei from denervated muscles. Dot size represents the percentage of nuclei expressing a gene. The color intensity of the dot indicates the expression level. Kruskal-Wallis test, ***p <0.001 compared to normal. (D) Heatmap showing changes in the normalized scores of L-R interactions between type I myofibers and other cells in normal and denervated condition. The interaction score is determined by the expression level of Fgf1 ligand in myofibers and expression level of receptors in receiver cells. The columns represent the receiver cell types and the rows represent predicted Fgf1 and receptor pairs. The color scale represents the score of interaction; a higher score indicates greater interaction between cells. (E) Heatmap of ligand-receptor interaction scores. Results show that the interactions between ligands (Fgf13 or Vegfa in type II myofibers) and their receptors in other resident cells.

Since Fgf1 and Vegfa are critical regulators of skeletal muscle homeostasis and vascular remodeling [35, 36], we evaluated the interactions mediated by these ligands. In normal muscles, we observed high scores for L-R pairs involving Fgf1 and its receptors. Upon denervation, these scores sharply decreased, implying abundant communications between type I fiber and other neighboring cells were blocked (Figure 9D). For example, the interaction between myofiber and MuSCs through Fgf1/Fgfr4 was weakened. Since Fgf1 stimulates the proliferation of MuSCs, this may suggest that decreased Fgf1 secretion from myofiber ultimately diminishes the proliferation and differentiation of MuSCs in denervated muscles. In addition, Vegfa-mediated communications between type II myofiber and resident cells, such as the Vegfa/Kdr interaction pair, were also restricted after denervation (Figure 9E). Attenuated Vegfa/Kdr signaling has been linked to denervation-induced vascular remodeling [37]. Regarding the ligand Fgf13, it was reportedly involved in neuron migration and stabilization of microtubules [38]. However, Fgf13 is an intracellular protein and has no identified interactions with Fgf receptors [39], thus its role in muscle atrophy needs further investigation. Taken together, these findings indicate that denervation perturbs the signals from myofibers, inducing different responses in various types of resident cells and consequently contributing to muscle atrophy.

## Discussion

Skeletal muscle exhibits tremendous plasticity to adapt to changes in the environment, a process involving protein synthesis and degradation, metabolic reprogramming, and tissue remodeling. This capability is fully manifested in denervated muscle. Type I myofibers, which support endurance activity, are known to be profoundly affected by denervation [23]; our snRNA-seq not only supported this conclusion but also evidenced even more robust transcription profile changes in type II fibers, which manifested as a dramatic shifting of the type IIb2 trajectory upon denervation (Figure 2). Moreover, the proportion of type IIb2 myonuclei was increased in conjunction with decreased proportions of type IIa/x myonuclei. Existing evidence supports that myonuclei are not lost during muscle atrophy [40], thus the observed proportion changes indicate the occurrence of myonuclear type transitions (such as type IIa to type IIb2). Thus, we propose that such transitions could be the root mechanism underpinning muscle plasticity.

We characterized the metabolic landscape of myofibers and their adaptive responses to denervation. Consistent with previous studies [41], in normal muscles, we found type I and type IIa myonuclei to be characterized by higher activities in oxidative phosphorylation, the TCA cycle, and lipid metabolism, while type IIx and type IIb myonuclei primarily rely on glycolysis and gluconeogenesis. In response to denervation, major metabolic pathways were largely downregulated, which raises the possibility that myofibers adapt to the unfavorable environment by reducing their overall energy demands and entering an inactive state. The observation that denervation suppresses the expression of both oxidative and glycolytic metabolism related genes adds an amendment to existing notion that denervation induces an oxidative to glycolytic fiber type switch. This could be due to our prediction being based on gene expression profiles, which do not fully reflect the actual enzyme activities. Overall, our results indicate that the decrease in metabolism is an important aspect of skeletal muscle adaptive responses upon denervation, which in turn could contribute to muscle atrophy [42].

We further ascertained the gene regulatory networks which drive cell type-specific transcriptomes under normal and denervated conditions. Muscle atrophy is mediated by a transcription-dependent program involving the upregulation or downregulation of a group of atrophy-related genes, such as Trim63, Fbxo32 [42]. Indeed, the activities of the Foxo3, Runx1, and Mef2a regulons were robustly increased in the type IIb1/2 myonuclei of denervated muscles (Figure S3A), consistent with previous reports that these TFs are important regulators of atrophy-related genes expression and muscle atrophy [43]. The effects of other regulons in the pathogenesis of muscle atrophy remain unidentified. Robo2 and Meis1 are known to be essential for embryonic skeletal muscle development [22, 44]; activation of their regulons in denervation implicates activation of an early developmental program in the pathogenesis of muscle atrophy. Bhlhe40 has been shown to control oxidative metabolism in myogenic cells [45], while *Elk4* has been reported to be a target gene downstream of MAPKs. Zfp369 belongs to the krueppel C2H2-type zinc-finger protein family and has been proposed to act as a transcription repressor [46]. Validation of these TF-related regulons as promoting atrophy-related genes expression and consequent muscle atrophy would add novel members to the atrophy paradigm.

Our results provide detailed insights into the transcriptome repertoire of muscle cells, including both type I and type II myofibers, and muscle-resident cells in the scenario of muscle atrophy. The results also highlight potential cellular and molecular targets for combating muscle atrophy induced by denervation or, perhaps, other related catabolic conditions.

---

**Figure S1.**
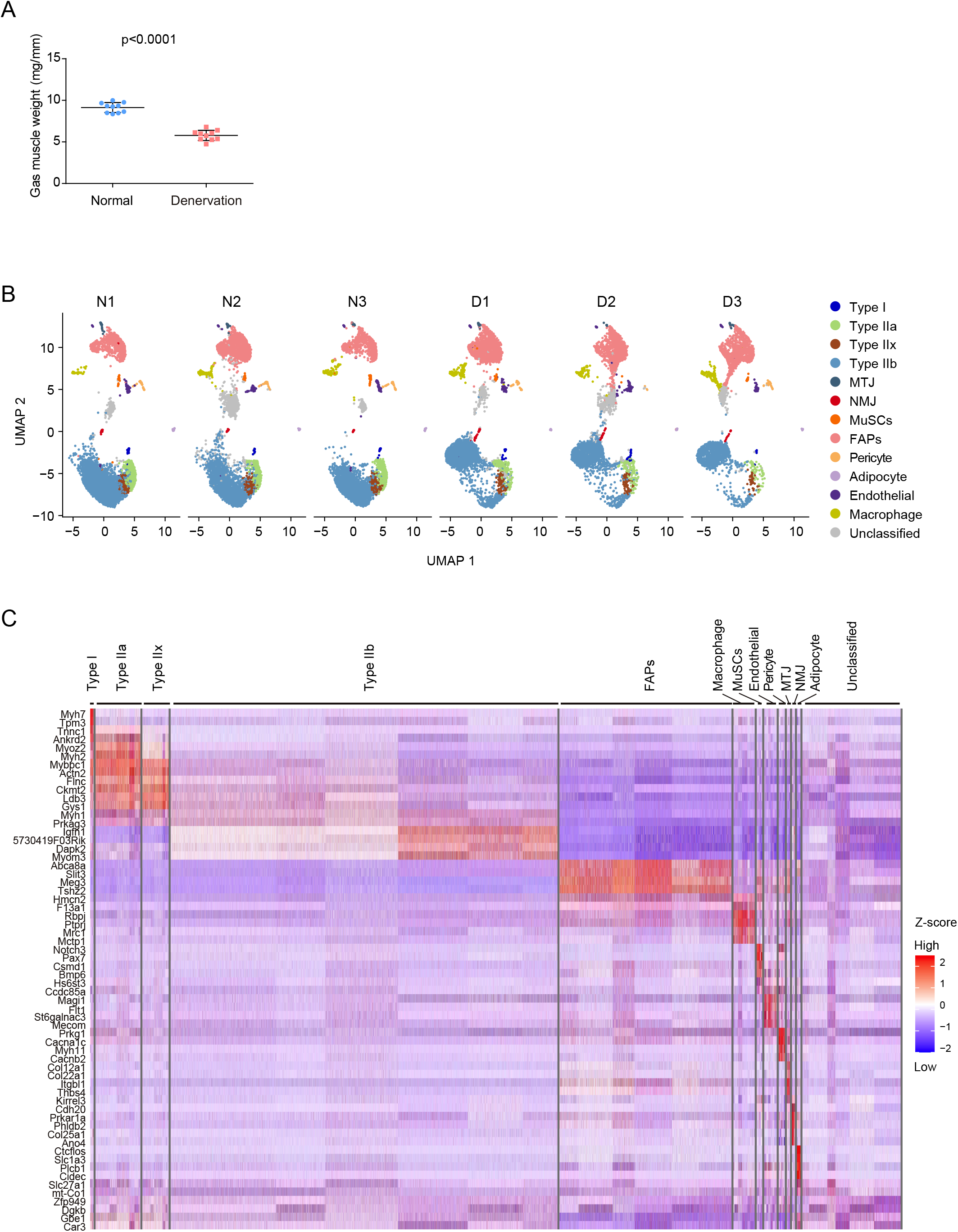
UMAP of clustering on sample sources and signature genes identification. (A) Gastrocnemius (Gas) muscle weight (normalized to tibia length) in normal and denervated conditions (n=10). Data are presented as mean ± SEM. Student’s t-test. (B) Visualization of the nuclei clusters on UMAP split by sample sources. All clusters are contributed by each of the normal and denervated muscle samples. Each cluster is color coded and labeled with sample types, N represents normal sample and D represents denervated sample, followed by sample number (N1-3 and D1-3). (C) Heatmap showing the top 5 signature genes of each cluster, identified by Seurat “FindAllMarkers” function across the whole dataset (logfc>0.25, min. pct>0.25).

**Figure S2.**
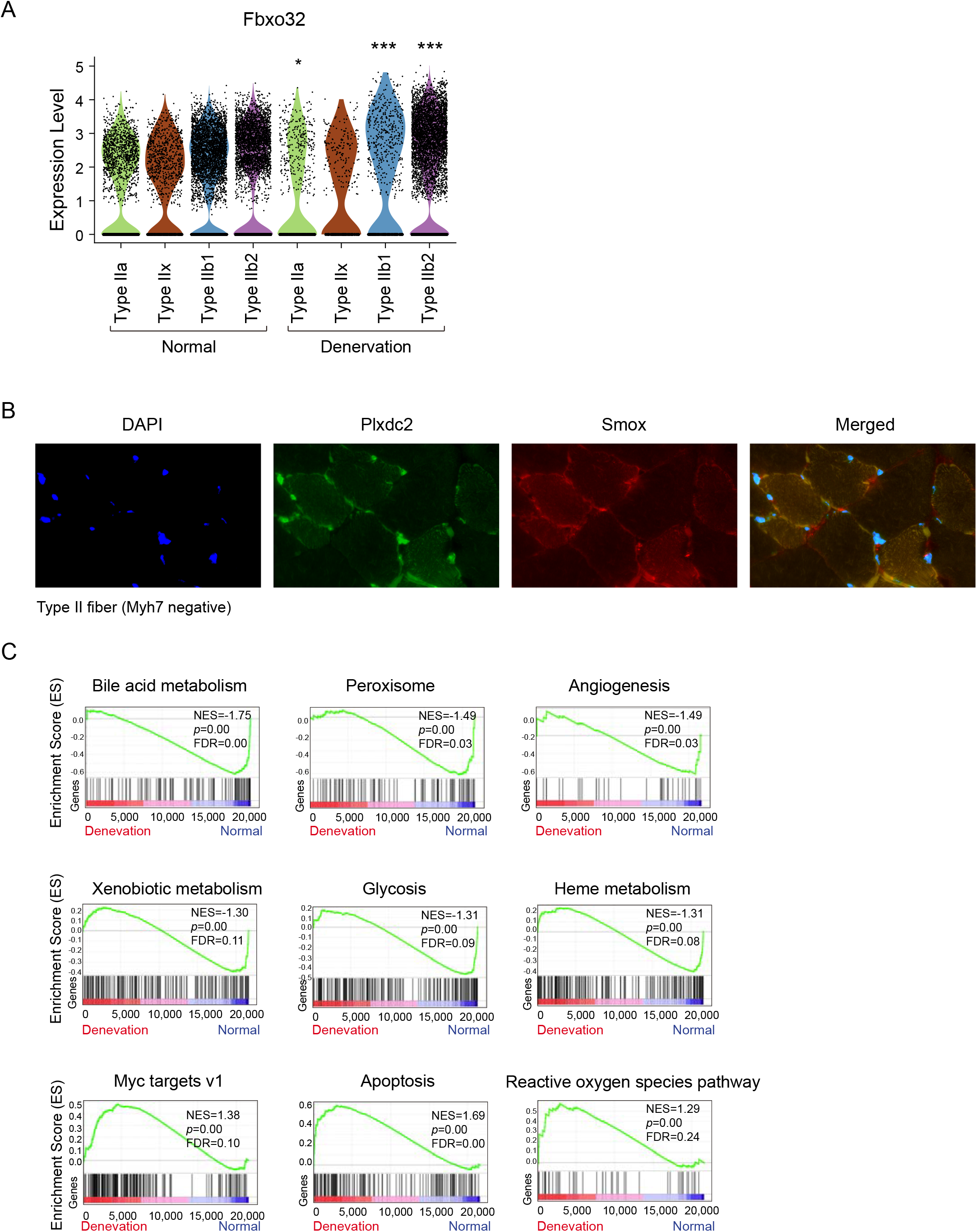
Transcriptional changes in type IIb2 myonuclei respond to denervation. (A) Violin plot showing significantly increased expression of Fbxo32 in denervated type IIa and type IIb myonuclei compared with normal. Wilcoxon rank-sum test, significance level: \**p* <0.05, \*\*\**p* <0.001. (B) Immunofluorescence staining of Plxdc2 (green, signature gene for type IIb2 nuclei) and Smox (red, signature gene for type IIb1 nuclei) in Myh7-negative type II fibers. Nuclei were labeled with DAPI (blue). (C) GSEA plots showing enrichment scores (ES) of the enriched hallmark gene sets in type IIb2 myonuclei. NES: normalized enrichment score; FDR: false discovery rate.

**Figure S3.**
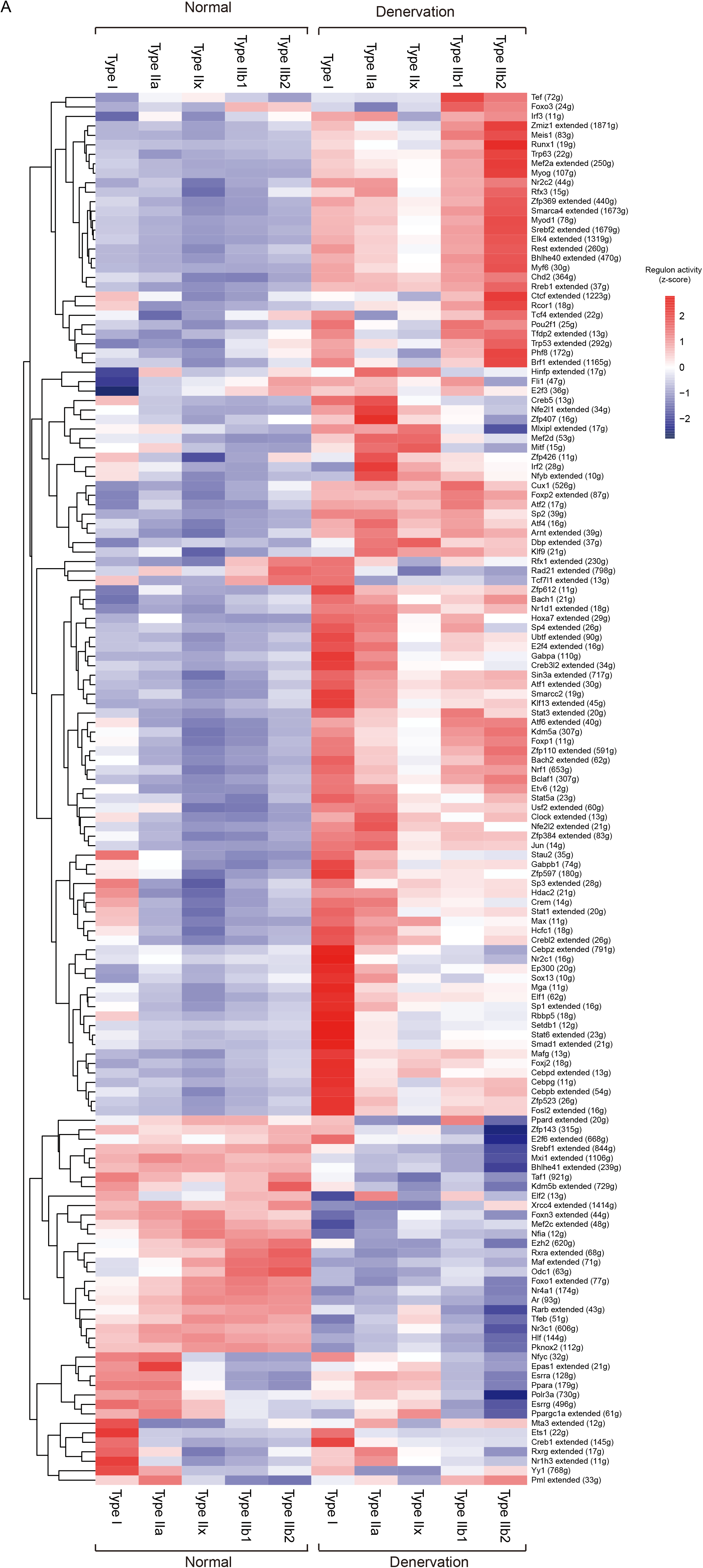
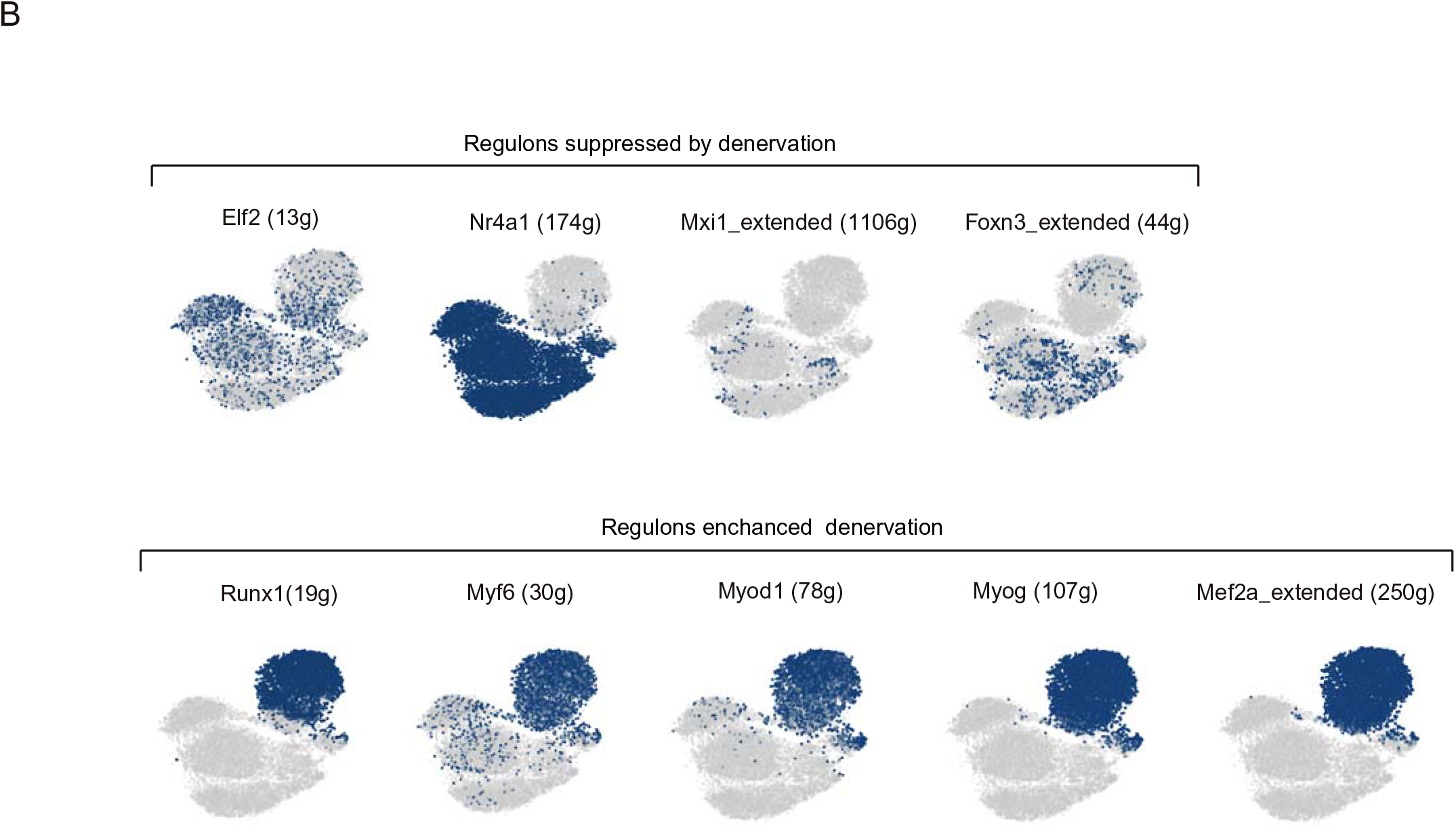
Regulons modulating type I and II myonuclei in normal and denervated muscles. (A) Heatmap showing the activity scores of total regulons (147) driving different myonuclei subtypes. The color scale represents the normalized scores of activities: red denotes high level of activity while blue indicates low level of activity. The number in parentheses represents the count of downstream target genes in corresponding regulons. (B) t-SNE on the binary regulon activity matrix showing that the regulon activity in type IIb2 myonuclei was suppressed (upper panel) or enhanced (lower panel) by denervation.

**Figure S4.**
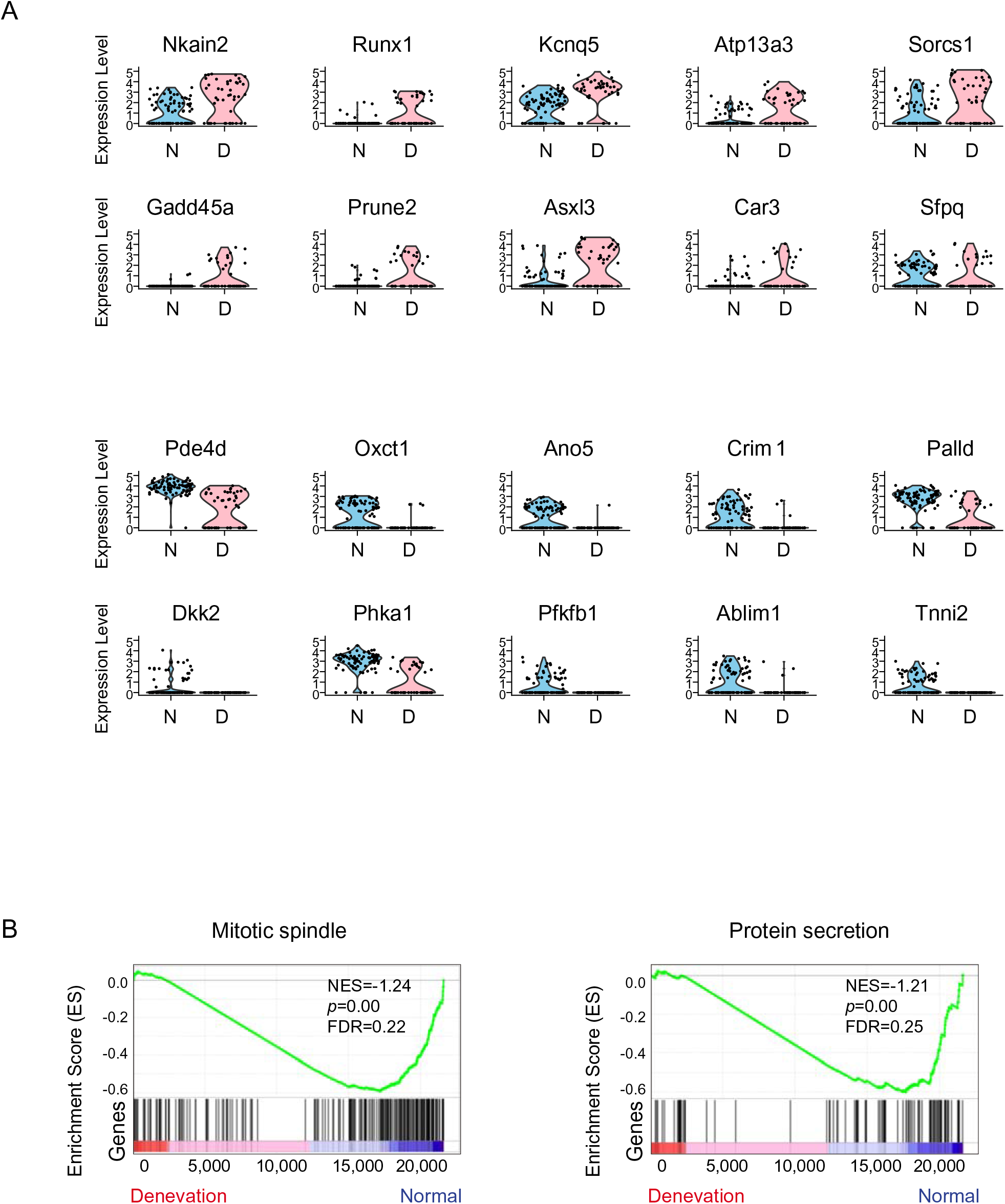
Transcriptomic changes of type I myonuclei in response to denervation. (A) Violin plot showing the significantly changed genes (logfc>0.25, min. pct>0.25, *p* < 0.001) between normal (blue) and denervated (pink) type I myonuclei. N: normal; D: denervation. Upper panel: up-regulated DEGs; Lower panel: down-regulated DEGs. (B) GSEA plots showing the enrichment score (ES) of significant enriched hallmark gene sets in type I myonuclei. NES: normalized enrichment score; FDR: false discovery rate.

**Figure S5.**
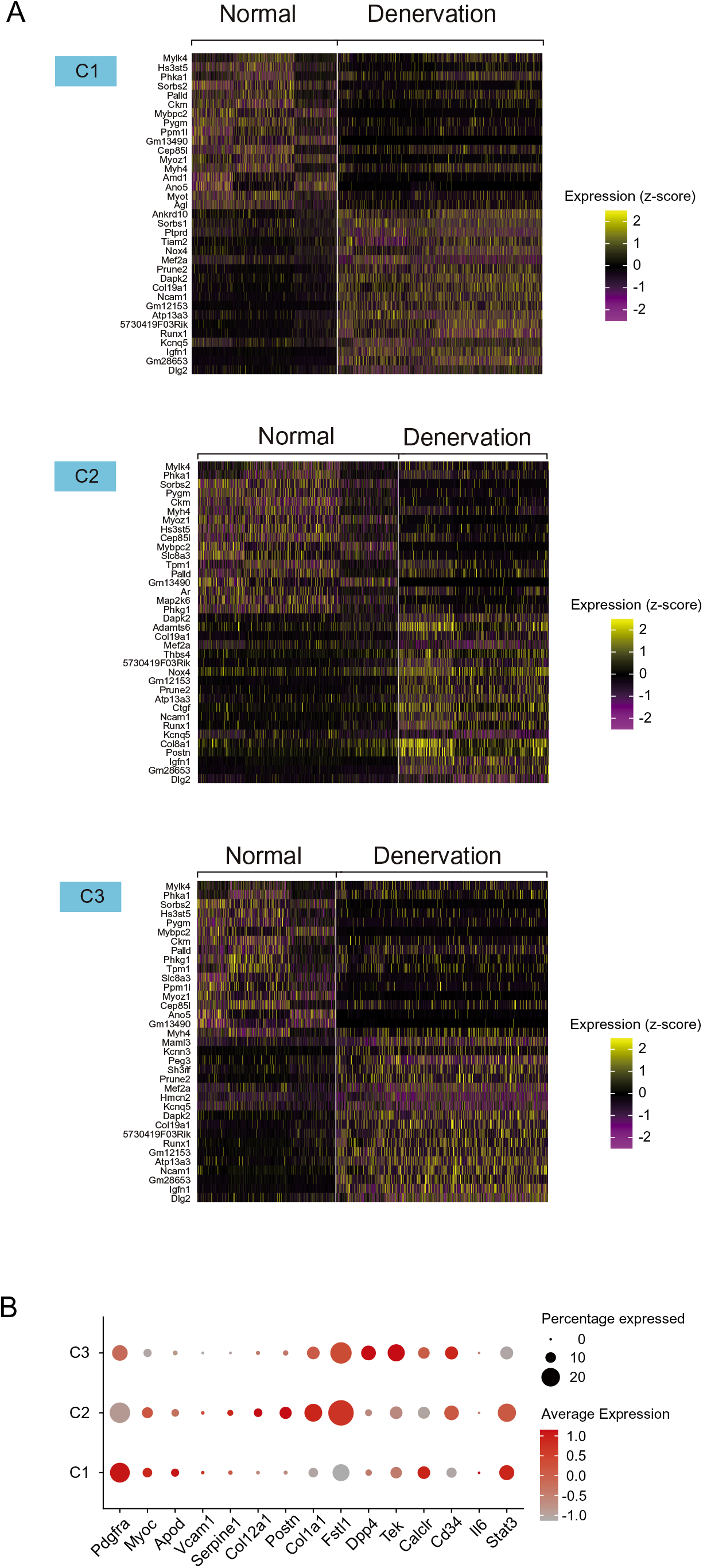
Denervation induced changes in FAPs. (A) Heatmap displaying the relative expression level of the top 20 DEGs between normal and denervation condition in 3 identified FAPs subtypes. Upper panel: DEGs of C1 subtype. Middle panel: DEGs of C2 subtype. Lower panel: DEGs of C3 subtype. The color scale represents the normalized expression values. (B) Dot plot displaying the average expression level of previous known FAPs identities in 3 FAPs subtypes. Dot size represents the percentage of cells expressing a gene within the subtype. The color intensity of the dot indicates gene expression level and darker color indicates higher expression level.

**Figure S6.**
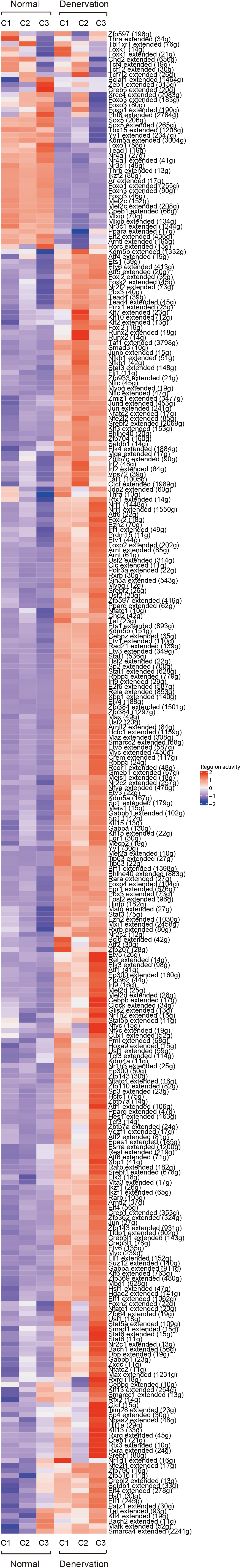
Corresponding changes of regulon activities in FAPs. (A) Heatmap showing the changes in activity scores of regulons (297) in FAPs subtypes between normal and denervation. The color scale represents the normalized scores of regulon activity: red indicates high level of activity and blue indicates low level of activity. The number in parentheses represents the count of downstream target genes in corresponding regulons.

**Figure S7.**
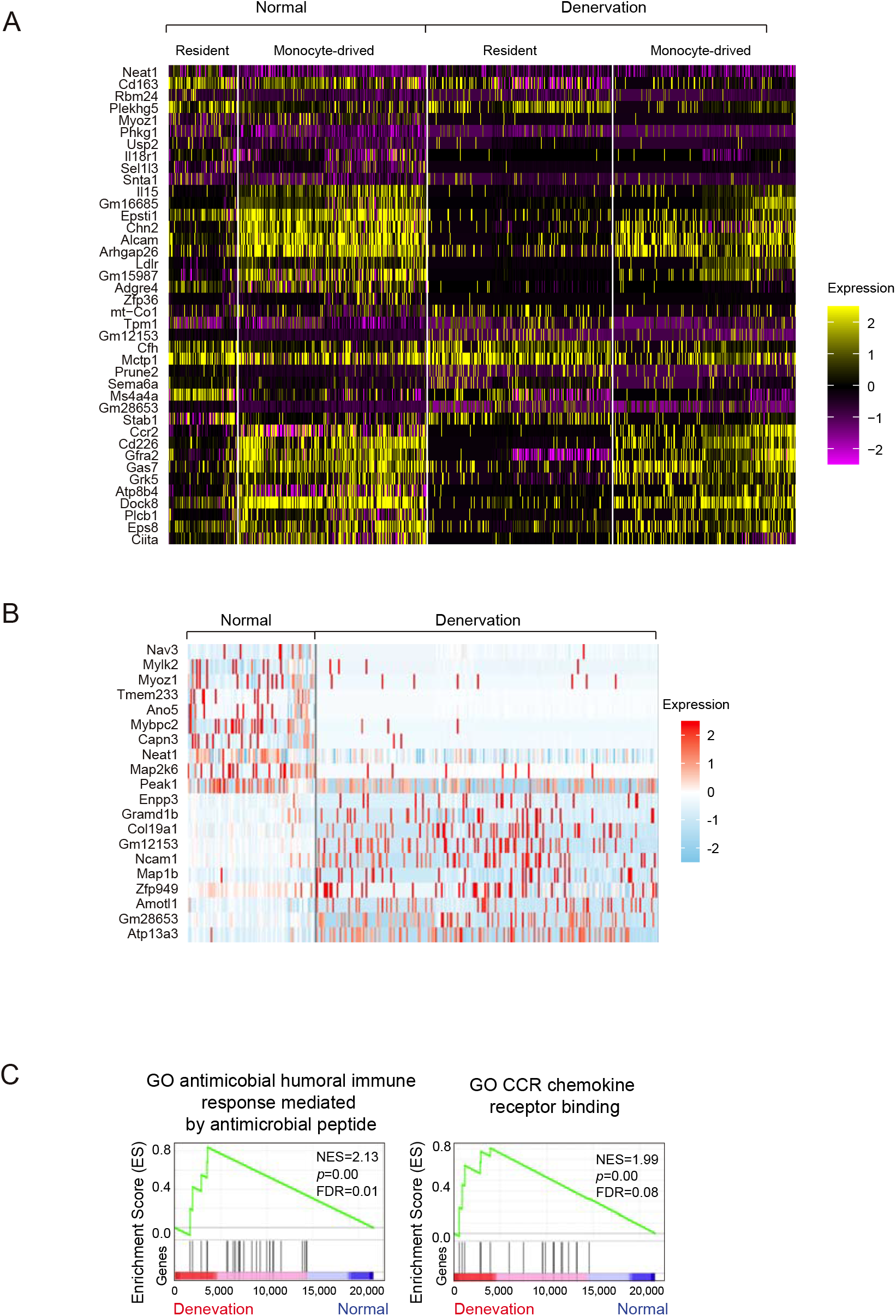
The macrophages in denervated muscles. (A) Signature genes of skeletal muscle-resident macrophages and monocyte-derived macrophages in skeletal muscle. (B) The expression levels of top10 DEGs in resident macrophages between normal and denervation. (C) Significant enriched GO gene sets in resident macrophages from denervated muscles. GO, Gene Ontology; NES: normalized enrichment score; FDR: false discovery rate.

**Figure S8.**
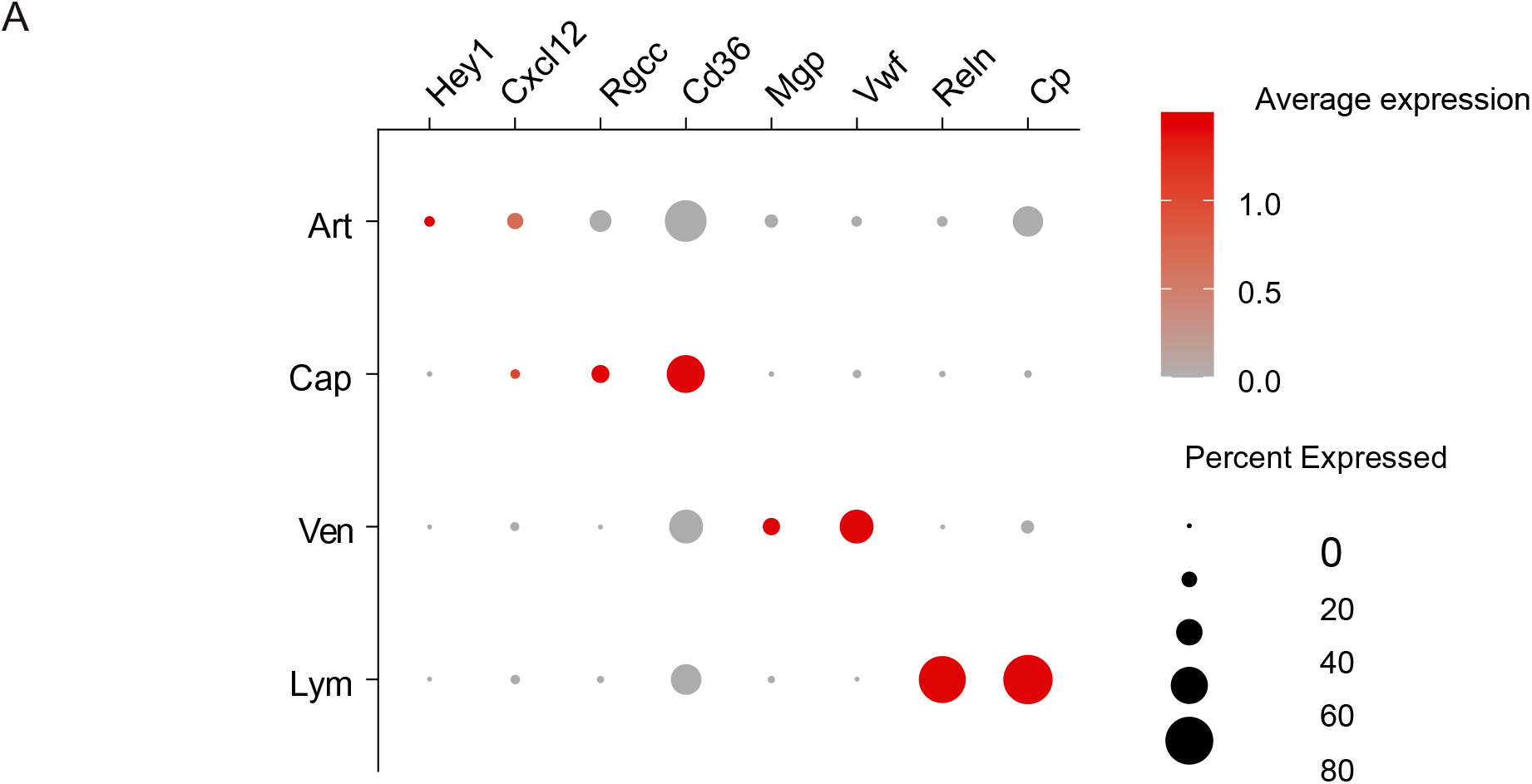
The cell identities for endothelial cells (ECs) from skeletal muscles. (A) Dot plot displaying the cell identities of 4 ECs subtypes in normal muscles. Dot size represents the percentage of cells expressing a gene within the subtype. Dot size represents the percentage of cells expressing a gene within the subtype. The color intensity of dot indicates gene expression level and darker indicates higher expression level.

**Figure S9.**
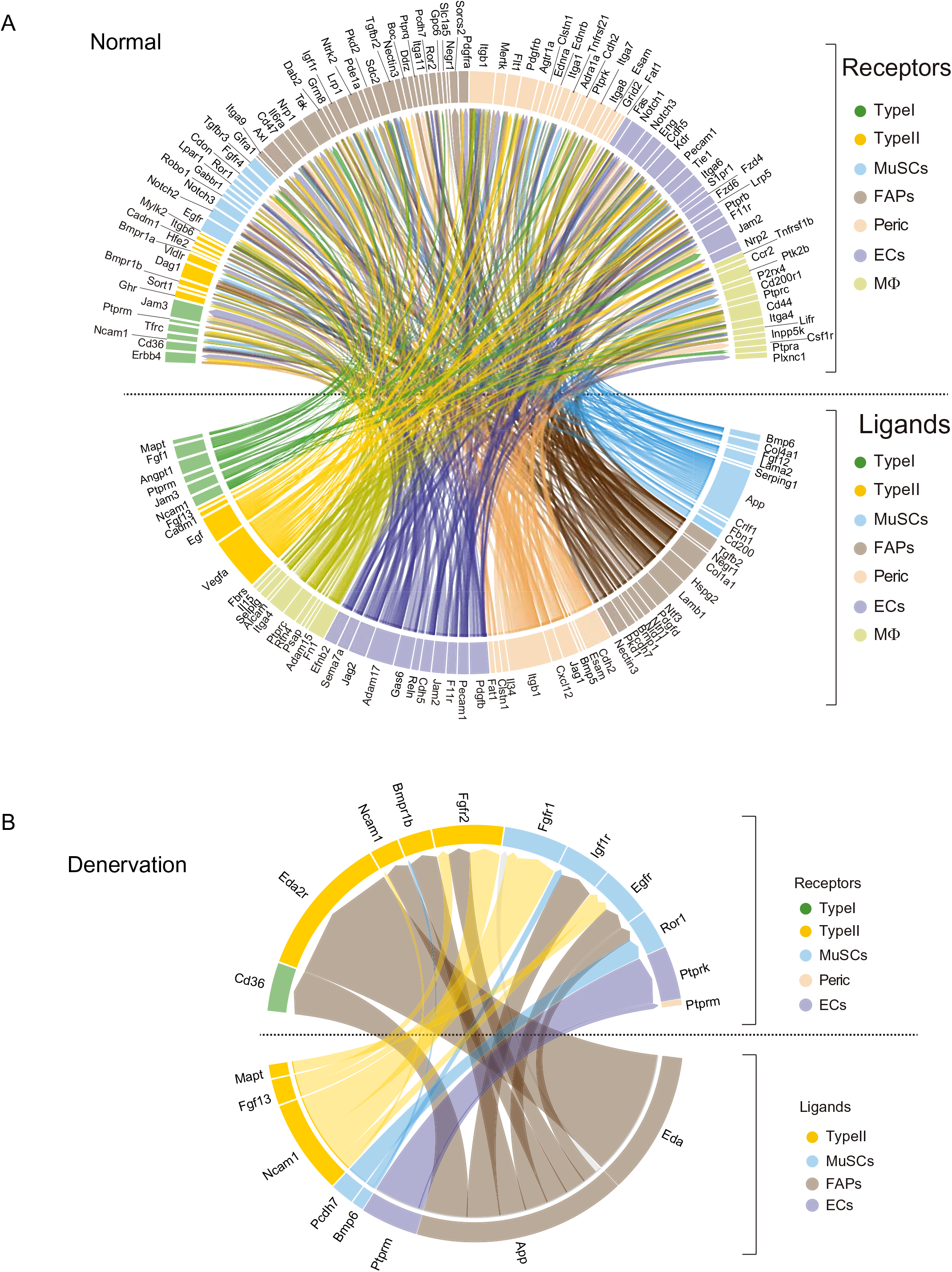
Cell -cell communications in skeletal muscle mediated by ligand-receptor (L-R) interactions. (A) Chrod plot displaying potential L-R interaction pairs between cells in normal muscles. A ligand-receptor (L-R) interaction model was used to calculate the scores for L-R pairs based on the expression levels of ligands and receptors. All types of cells function as either sender (ligand expressing) or receiver (receptor expressing) cells. Only differentially expressed ligands and receptors (logfc>0.25, min. pct>0.25) are considered for further analysis. The detected ligands are listed below the dash line, and receptors are listed above the dash line. Each cell type is color-coded. (B) Chrod plot displaying potential L-R interaction pairs between cells in denervated muscles.

